# An *N,S*-acetylated L-cysteine-cysteamine conjugate hinders pyocyanin redox cycling to weaken *Pseudomonas aeruginosa* biofilm and dampens LPS-driven acute pulmonary inflammation

**DOI:** 10.64898/2026.05.13.724891

**Authors:** M Bruschi, S Masini, F Palma, Y Xiaoqiu, CL Braga, M Gregori, C Bucci, F Bartoccini, M Menotta, E Manuali, L Minelli, D Ligi, F Mannello, F Monittola, C Zara, C Di Pietro, R Crinelli, G Brandi, G Piersanti, EM Bruscia, GF Schiavano, A Fraternale

**Affiliations:** Department of Biomolecular Sciences, University of Urbino Carlo Bo, Urbino, PU, Italy; Department of Pediatrics, Yale School of Medicine, New Haven, CT, USA; Experimental Zooprophylactic Institute [Umbria and Marche] “Togo Rosati”, Perugia, PG, Italy; Department of Humanities, University of Urbino Carlo Bo, Urbino, PU, Italy

**Keywords:** *Pseudomonas aeruginosa*, redox, biofilm, pyocyanin, ROS, thiols

## Abstract

The persistence of *P. aeruginosa* infections is largely driven by the secretion of several factors during invasion, including the redox-active phenazine pyocyanin (PYO), which promotes biofilm formation and oxidative stress. Biofilms contribute to chronic infections and antibiotic resistance, limiting the efficacy of conventional therapies. We found that a synthetic compound, I-152, a conjugate of N-acetyl-L-cysteine (NAC) and S-acetylcysteamine (also known as S-acetyl-β-mercaptoethylamine; SMEA), effectively restored colistin susceptibility against *P. aeruginosa* by altering biofilm nanomechanical properties. These perturbations in matrix integrity were associated with I-152’s ability to hinder the phenazine redox cycle, shifting PYO to a reduced state as well as enabling S-conjugate formation. The compound decreased PYO accumulation in bacterial cultures and PYO-generated reactive oxygen species (ROS) in macrophage cells. Together with PYO, LPS is another driver of ROS-dependent inflammatory signaling in the host, which leads to an uncontrolled cytokine response and organ damage, especially in patients with cystic fibrosis. I-152 treatment downregulated the expression of LPS-induced inflammatory cytokines, i.e., IL-6 and TNF-α, in bone marrow-derived macrophages (BMDM) isolated from transgenic CFTR^-/-^ and CFTR^+/+^ mice. Consistently, I-152 partially counteracted the inflammatory response in the *P. aeruginosa* LPS-induced acute lung injury murine model. Taken together, these results support I-152 as an adjunctive treatment for *P. aeruginosa* respiratory infections through a dual mechanism: combating antimicrobial resistance in biofilms and dampening host inflammation in the respiratory system.

**Highlights:**

- I-152 potentiates colistin activity against *P. aeruginosa* by compromising the biofilm surface
- I-152 rewires the pyocyanin (PYO) redox state and forms covalent adducts with it
- PYO accumulation and PYO-induced ROS generation in macrophages is impaired by I-152
- *Ex vivo,* I-152 dampens excessive pro-inflammatory response to *P. aeruginosa* LPS in CFTR^-^/^-^ and CFTR^+^/^+^ BM-derived macrophages
- I-152 (140 mg/Kg) attenuates LPS-driven inflammation and lung damage in CFTR^+^/^+^ mice

## 1. Introduction

*Pseudomonas aeruginosa* (*P. aeruginosa*) is a ubiquitous, opportunistic pathogen and has become one of the most common causes of nosocomial bacterial infections since it is widely found on medical devices (e.g., ventilators). The most severe *Pseudomonas* infections include respiratory infections in patients with cystic fibrosis (CF) and systemic bloodstream infections that have disseminated from burn wounds or pneumonia [1,2]. One complication, often encountered during treatment of *P. aeruginosa* infections, is the development of antibiotic resistance in the bacteria. *P. aeruginosa* has an extraordinary capacity to gain resistance via multiple mechanisms, even concurrently, resulting in the ability to withstand nearly all available antibiotics.

An additional pathogenetic factor leading to therapeutic failure is the complex structure of the *P. aeruginosa* biofilm [3]. The biofilm matrix primarily comprises exopolysaccharides, extracellular DNA (eDNA), and matrix proteins, which play essential roles in structural maintenance, antibiotic resistance, and the development of chronic infections that are difficult to eradicate. [3]. *P. aeruginosa* can adapt to the environment in hosts by secreting a variety of virulence factors that have been implicated in contributing to the maintenance of hyper-inflammatory conditions and successful infection [4]. Among these virulence factors, pyocyanin (PYO) is a redox-active phenazine, membrane-permeable, that by interacting with intracellular pools of NAD(P)H and glutathione (GSH), depletes the mammalian cells of these compounds, and produces reactive oxygen species (ROS), especially O_2_^−^ and H_2_O_2_, resulting in oxidative damage to different cellular components [5–7]. The ability of PYO to induce redox alteration and increase oxidative stress appears central to its diverse detrimental effects on host cells. [7] Additionally, *P. aeruginosa* overcomes oxygen limitation within the biofilm by secreting soluble redox-active phenazines, such as PYO, which act as electron shuttles to distal oxygen. Therefore, it has been recently reported that biofilm formation can be inhibited electrochemically by controlling the redox state of PYO [8].

Due to these circumstances*, P. aeruginosa*-infected airways form a dysregulated, hyperinflammatory environment, with elevated levels of inflammatory cytokines, mucus overproduction, and increased numbers of infiltrated and activated neutrophils, which significantly contribute to disease severity [9]. Lipopolysaccharide (LPS) is another prominent pathogenic factor that mediates both bacterial virulence and host responses [10]. Among the different sites of infection, the lungs are one of the first interfaces to encounter *P. aeruginosa* bacteria and their LPS, and they elicit the initial host responses. Lungs are highly exposed to both endogenous oxidants, *e.g.*, ROS produced by activated phagocytes, and exogenous oxidants, both inhaled and those produced by pathogens [6,7,10], so a reducing environment composed of enzymatic and non-enzymatic antioxidants could prevent an exaggerated inflammatory response. This aspect is crucial in CF lungs, where *P. aeruginosa* accelerates the decline in pulmonary function. In fact, they are characterized by an oxidative environment, which alters ion transport, including the already impaired cystic fibrosis transmembrane conductance regulator (CFTR) [11]. Therefore, the redox state appears to play a pivotal role in *P. aeruginosa* infection and in an efficient/balanced host response, resulting in a valuable target for new therapeutic approaches. Low-molecular-weight (LMW) thiols, which contain reducing sulfhydryl groups, affect redox chemistry in bacteria and may also play a role in virulence and bacterial pathogenesis [12,13]. For example, we have previously reported that LMW thiols exerted bacteriostatic and bactericidal activity against *Mycobacterium avium* (*M. avium*) and modulated macrophage immune response towards the bacterium [14].

Other groups described how GSH neutralized oxidative stress mediated by PYO and how N-acetyl-L-cysteine (NAC) attenuated quorum-sensing-regulated phenotypes in *P. aeruginosa* PAO1 [15,16]. Recently, Liu and colleagues explored the use of reducing agents to inhibit biofilm formation and enhance biofilm susceptibility to antimicrobial treatments [17]. In this work, we report results regarding the antibacterial activity of a LMW compound, *i.e.*, I-152, a conjugate of NAC and S-acetyl-β-mercaptoethylamine (SMEA), against *P. aeruginosa*. We demonstrate that I-152 inhibited PYO accumulation in bacterial cultures, both interfering with its redox cycling and by directly reacting with it, leading to enhanced colistin activity against *P. aeruginosa* and modified biofilm physical characteristics. Moreover, the inflammatory response to *P. aeruginosa* LPS insult was assessed *ex vivo* in bone marrow monocyte-derived macrophages (BMDM) coming from wild-type CFTR^+/+^ (WT) and CFTR^-/-^ (CF) mice and *in vivo* in WT mice. I-152 treatment affected cytokine responses in both experimental models and attenuated LPS-induced murine lung injury. Hence, redox-based strategies could represent an effective approach for biofilm management and protection against lung damage by interfering with PYO redox status and modulating cytokine responses.

## 2. Materials and Methods

### 2.1 I-152 synthesis

The compound was synthesized following previously established protocols (compound 6) [18]. In *in vitro* experiments, *P. aeruginosa* LPS (Sigma-Aldrich) was prepared as a 100× stock solution in PBS (phosphate-buffered saline) and used at a concentration of 10 μg/mL.

### 2.2 Bacterial strain and antimicrobial activity

The strain used in this study was *P. aeruginosa* ATCC 27853. To prepare bacterial broth cultures, bacteria from a single colony on a *Pseudomonas* agar plate were inoculated into 20 mL of Mueller-Hinton Broth (MHB, Liofilchem) and incubated overnight (16-18h) at 37°C to reach a cell concentration of approximately 5×10^^8^ CFU/mL.

Three mL of bacterial suspensions of *P. aeruginosa* (1×10^^6^ CFU/mL) in saline solution (0.85% NaCl) or MHB (1×10^^5^ CFU/mL), were treated in triplicate with different concentrations of I-152 (0.25 mM, 1 mM and 3 mM) and incubated at 37° C. Samples were taken at different time intervals for enumeration of viable *P. aeruginosa* (CFU/mL) on *Pseudomonas* selective agar (PAB, Oxoid).

### 2.3 *In vitro* quantitative assay for planktonic growth and biofilm formation

The overnight culture was diluted to about 1×10^^6^ CFU/mL in sterile MHB, and 500 µL were dispensed into each well of a sterile flat-bottom 48-well plate. Plates were incubated at 35°C under static conditions for different time frames, as depicted in the experimental protocol shown in Fig. 2. Non-adherent bacteria were removed by washing twice with 500 µl sterile PBS and used to assess planktonic growth, recording the optical density (OD) at 600 nm (OD600). The biofilm biomass was stained with 150 µl of 0.1% crystal violet per well for 20 min, and the surface-attached cells were quantified by solubilizing the dye in ethanol and measuring the OD at 570 nm (OD570) using a microplate reader (Tecan Spark® Multimode Microplate, Switzerland). Each data point was averaged from at least three replicate wells.

**Fig. 1.**
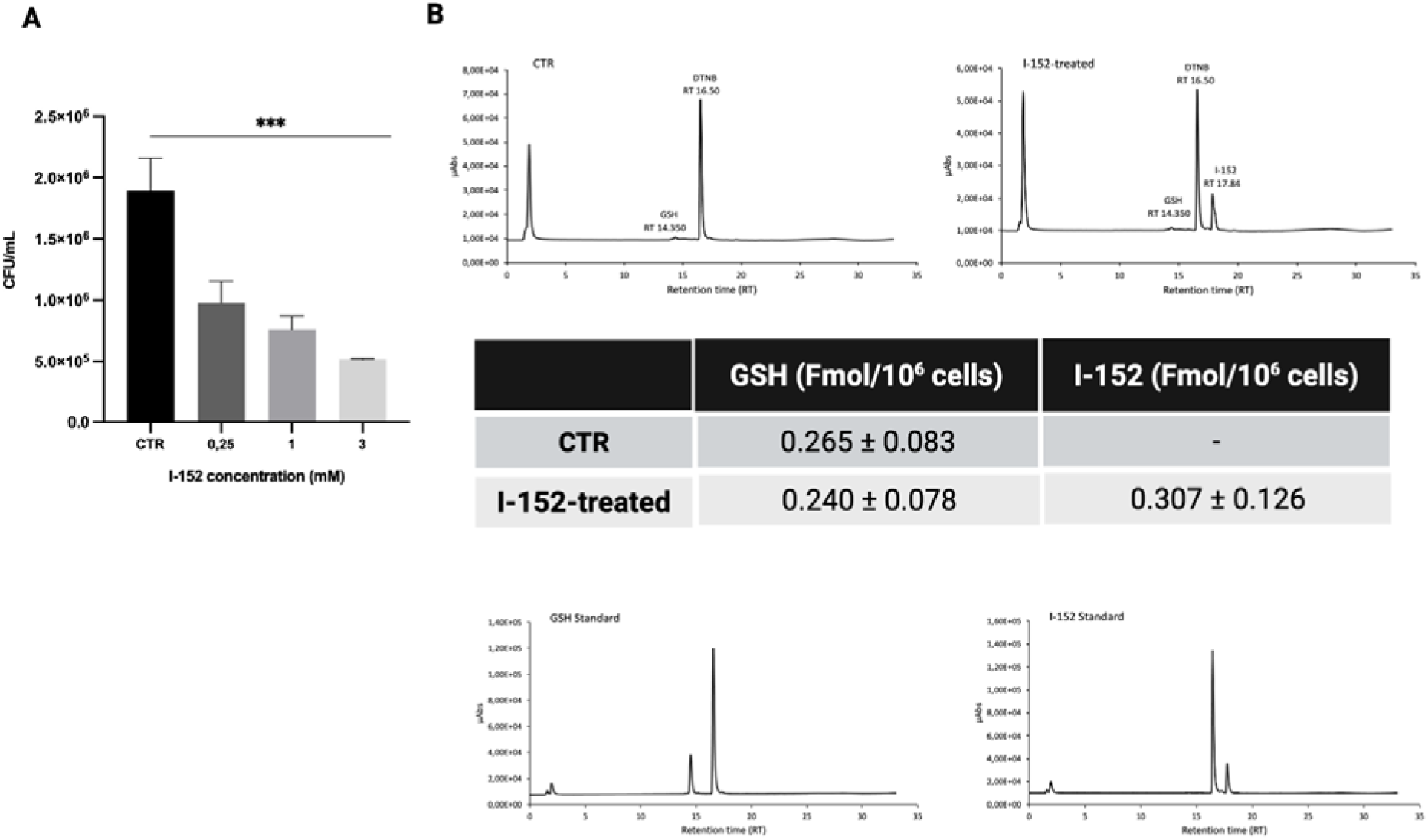
Antimicrobial activity of I-152 and its detection in *P. aeruginosa* suspensions. (**A**) *P. aeruginosa* suspensions in MHB were exposed to different concentrations of I-152, and after 1h-treatment, samples were plated on Pseudomonas isolation agar to determine the number of CFU/mL. Results are mean ± SD of 3 experiments. ***p<0.001. (**B**) I-152 was added to *P. aeruginosa* (1×10^^6^ cells/mL) at a concentration of 1 mM for 1h. After medium removal, the cells were lysed, and the proteins were precipitated. After centrifugation, reduced glutathione (GSH) and I-152 were determined in the supernatant by the HPLC method [19]. Above: an illustrative chromatogram of *P. aeruginosa* cells incubated for 1h with 1 mM I-152 (right) or not incubated (left), and in the table, quantification of GSH and I-152 in *P. aeruginosa* cells reported as femtomoles (Fmol)/10^^6^ cells; below: an illustrative chromatogram of GSH and I-152 standards (20 µM) used for quantification.

**Fig. 2.**
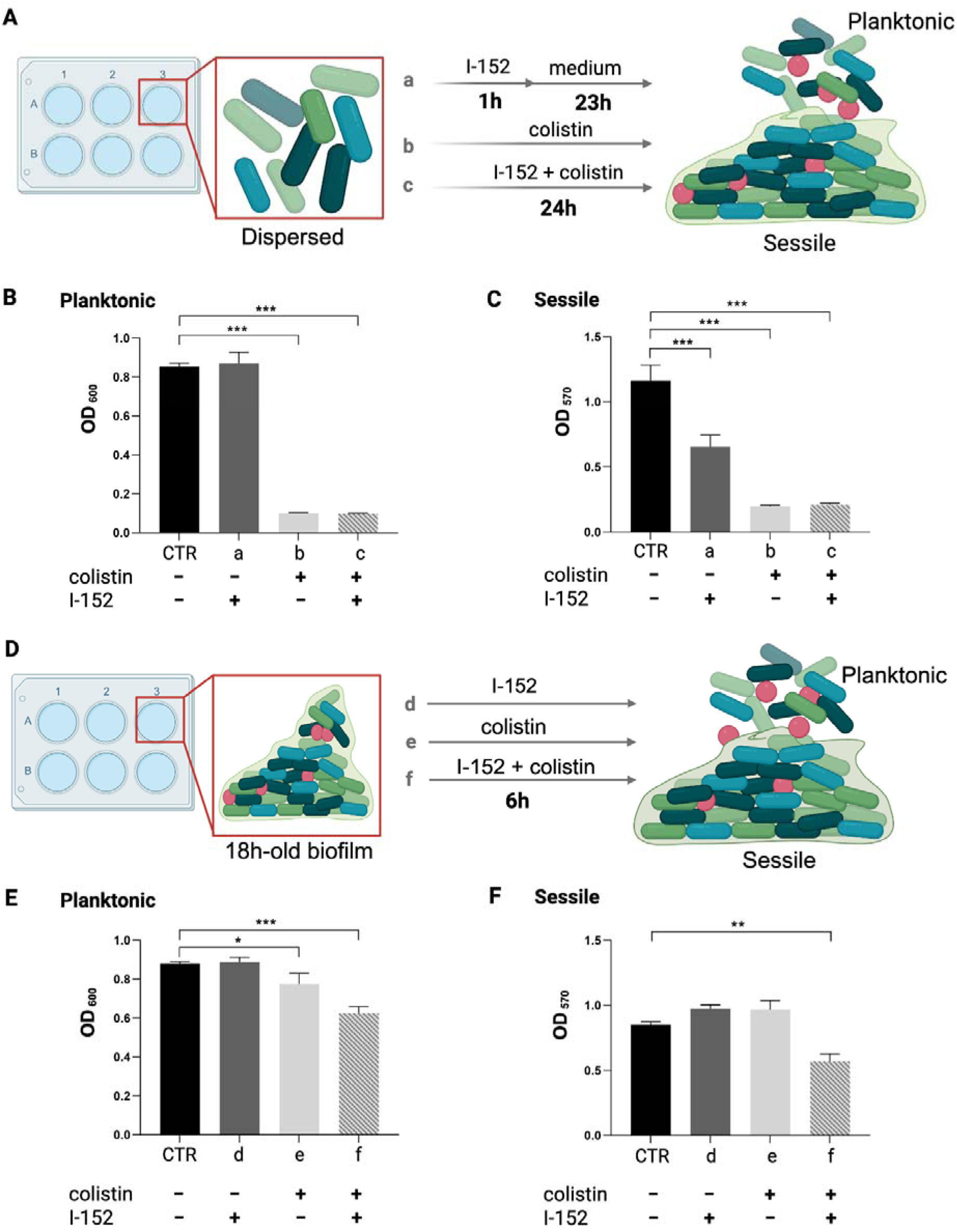
Effects of I-152, alone or in combination with colistin, on *P. aeruginosa* planktonic growth and biofilm formation. The treatment protocols are reported in (**A**) and (**D**). I-152 and colistin were administered at the concentrations of 3 mM and 4 mg/L, respectively. Specifically, the treatments were as follows: a) 1h I-152 and then the medium was added up to 24h; b) colistin was kept for 24h; c) I-152 was given in combination with colistin for 24h. After 18h-biofilm formation: e) I-152, f) colistin, and g) I-152 combined with colistin were given for 6h. (CTR) *P. aeruginosa* without any treatment. Planktonic growth (**B** and **E**) and biofilm biomass (**C** and **F**) were evaluated by absorbance readings in a microplate reader at 600 and 570 nm, respectively. Results are the mean ± SD of 3 experiments. *p<0.05, **p<0.01, ***p<0.001.

### 2.4 Determination of thiol species in *P. aeruginosa* cells

*P. aeruginosa* (5×10^^6^ bacteria/5 mL) were treated or not with 1 mM I-152 for 1h. At this time, the cells were pelleted by centrifugation at 100×g to remove the medium and washed twice with PBS. The cell pellet was lysed with 500 μL of lysis buffer (0.1% Triton X-100, 0.1 M Na_2_HPO_4_, 5 mM EDTA, pH 7.5) followed by 75 μL of 0.1 N HCl and 700 μL of precipitating solution [200 mM glacial metaphosphoric acid, 6 mM Na_2_EDTA, 5 M NaCl]. Before the addition of precipitating solution, the cell lysate was sonicated for 10 sec (Branson Sonifier, Model B-15, Branson Sonic Power, a SmithKline Company, power output 2.5 W/s) to achieve complete cell disruption; all procedures were carried out on ice. The samples were kept on ice for 10 min and then centrifuged at 12,000×g for 10 min at 4°C. Twenty-five % (v/v) Na_2_HPO_4_ 0.3 M and 4% (v/v) Ellman’s Reagent (5,5-dithio-bis-(2-nitrobenzoic acid) (DTNB) were added to the supernatants for thiol determination by HPLC through a BDS Hypersil™ C18 column (5 µm, 150×4.6 mm) (Thermo Scientific, USA). Elution conditions were previously described [19]. Detection was at 330 nm, and quantitative measurements were obtained using standard solutions and normalized to cell number.

### 2.5 Atomic force microscopy (AFM) analysis

AFM analysis was carried out with an XE-100 Atomic Force microscope (PARK Systems Inc., Suwon, South Korea). The microscope was equipped with a 50μm scanner controlled by the XEP 1.8.6 software. The XY stages and the Z scan were set in a closed-loop manner and in high-voltage mode. The speed scan was set between 0.2 Hz and 0.5 Hz. The instrument was set in Force Modulation Microscopy (FMM) achieved using an NSC36b tip (nominal force constant k= 2 N/m). The tip operational settings were determined by using the PARK Systems FMM standard sample. Local nanomechanical properties of the surface were evaluated by monitoring amplitude and phase signals of the scanned regions. Images were acquired randomly across the cell layers. For numerical comparisons, the FMM amplitude values of all pixels were analyzed using the Kruskal–Wallis test among groups. Additionally, FMM amplitude roughness parameters (Ra and Rq) were calculated by randomly collecting 2560 pixels (256 pixels x 10 lines) from each acquired image, followed by statistical comparison by using the Kruskal–Wallis (K-W) test.

### 2.6 PYO and I-152 ^1^H NMR analysis

PYO was synthesized as reported in the literature [20]. I-152 and its oxidized form (I-152_ox_) were synthesized as previously described [18].

^1^H NMR spectra were recorded on a 600 MHz spectrometer, using DMSO-*d6* as solvent. Chemical shifts (δ scale) are reported in ppm relative to the solvent peak.

Pyocyanin (1 mg, 0.0048 mmol) was dissolved in DMSO-*d6* (500 µL) in a standard 5 mm NMR tube, and the spectrum was recorded. Hantzsch ester (HE) (1.22 mg, 0.0048 mmol) was then added, and the spectrum was recorded 5 min, 20h, and 96h after the addition (Fig. 4A).

**Fig. 3.**
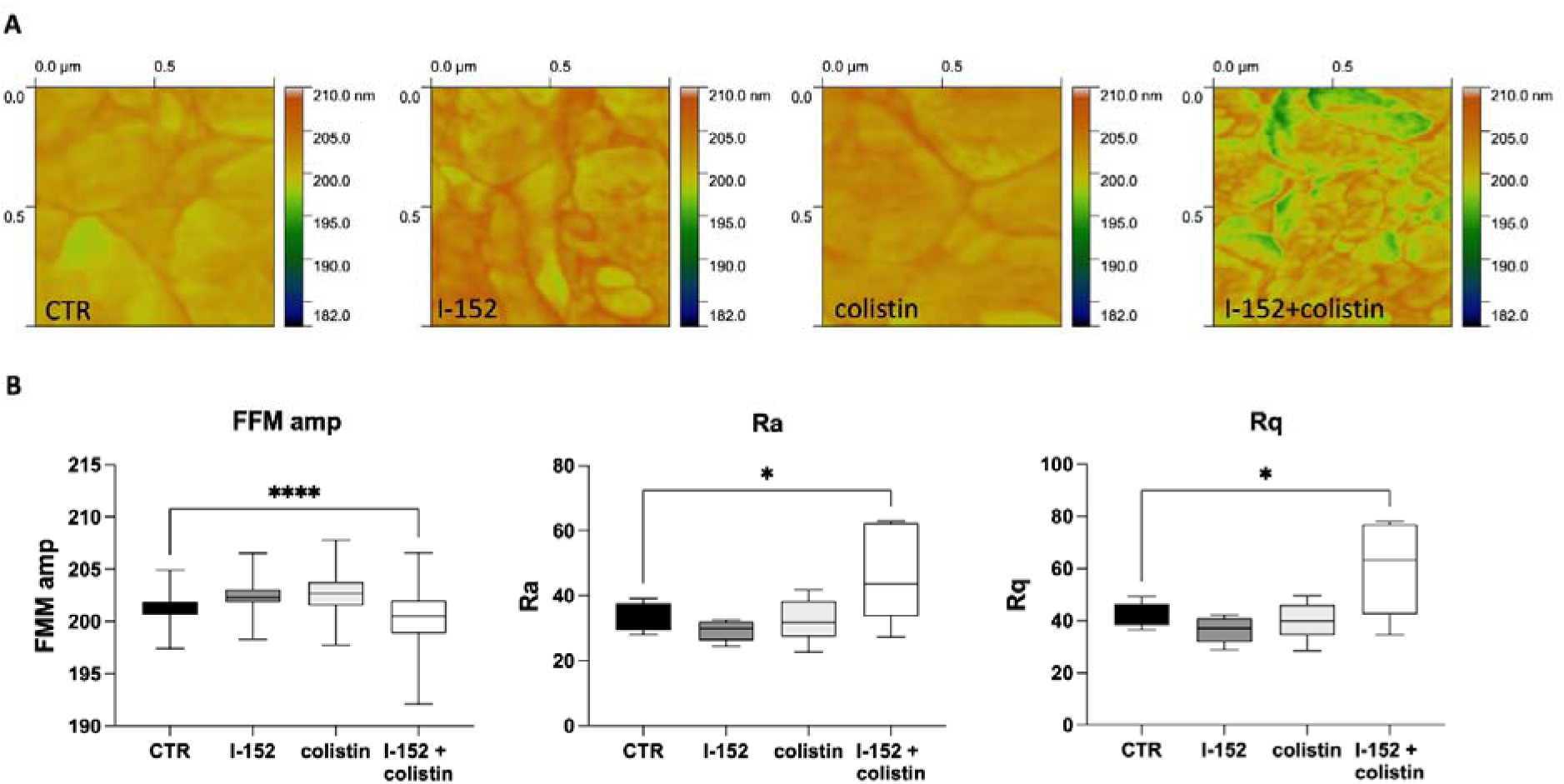
I-152 and colistin effects on the biofilm surface hallmarks. Nanomechanical features were analyzed after 18h of biofilm formation via AFM. Four experimental conditions were tested: CTR, *P. aeruginosa* which did not encounter any treatment up to 24h; colistin, *P. aeruginosa* received 4 mg/L on 18h-pre-formed biofilm for the next 6h; 3) I-152, *P. aeruginosa* received 3 mM I-152 on 18h-pre-formed biofilm for the next 6h; 4) I-152 + colistin, *P. aeruginosa* received 3 mM I-152 together with 4 mg/L colistin on 18h-pre-formed biofilm for the next 6h. (**A**) Representative AFM 3D topography images of *P. aeruginosa* biofilm surfaces (1 μm^2^) with tapping mode. (**B**) Box and whisker plots representing surface features, such as FFM amplitude, amplitude Roughness average (Ra) and amplitude Root mean square roughness (Rq), come from three independent experiments. *p < 0.05; ****p< 0.0001 K-W test followed by Dunn’s multiple comparisons.

**Fig. 4.**
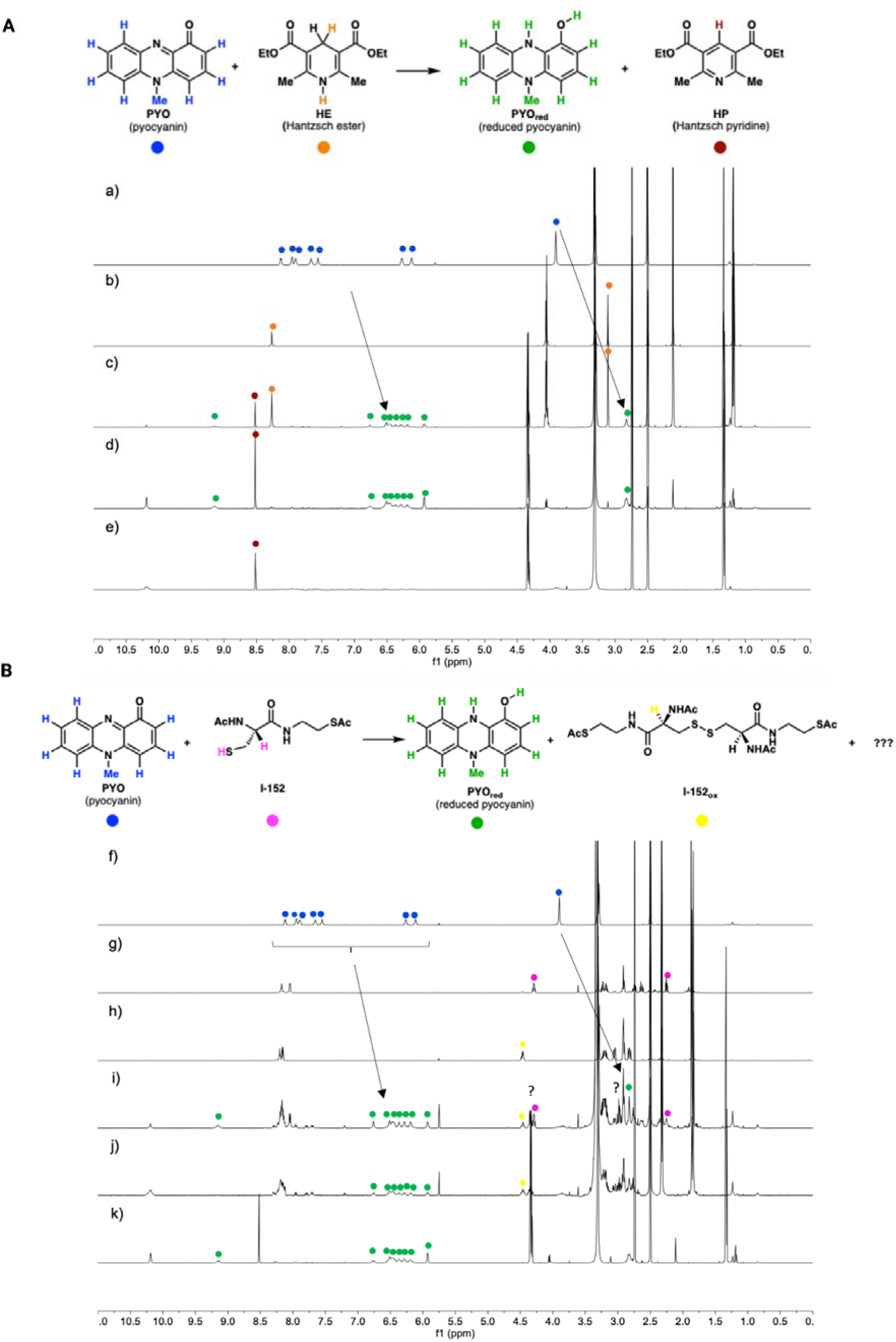
Chemical reaction of PYO with Hantzsch ester or I-152. (**A**) ^1^H NMR spectra inDMSO-*d_6_* of (a) PYO; (b) Hantzsch ester; (c) PYO (1 equiv) and HE (1 equiv) at the beginning of the reaction (5 min time of acquisition); (d) PYO (1 equiv) and HE (1 equiv), 20h; (e) PYO (1 equiv) and HE (1 equiv), 96h. (**B**) ^1^H NMR spectra in DMSO-d_6_ of (f) PYO; (g) I-152; (h) I-152_ox_; (i) PYO (1 equiv) and I-152 (1 equiv) at the beginning of the reaction (5 min time of acquisition); (j) PYO (1 equiv) and I-152 (1 equiv), 20h; (k) PYO (1 equiv) and I-152 (1 equiv), 96h. The reactions were carried out at RT.

Pyocyanin (1 mg, 0.0048 mmol) was dissolved in DMSO-*d6* (500 µL) in a standard 5 mm NMR tube, and the spectrum was recorded. Hantzsch ester (HE) (1.22 mg, 0.0048 mmol) was then added, and the spectrum was recorded 20h after addition. The solution was transferred to a vial and stirred at room temperature for 6h in an open vessel, and the spectrum was recorded (Fig. S1).

Pyocyanin (1 mg, 0.0048 mmol) was dissolved in DMSO-*d6* (500 µL) in a standard 5 mm NMR tube, and the spectrum was recorded. I-152 (1.27 mg, 0.0048 mmol) was then added, and the spectrum was recorded 5 min, 20h after addition (Fig. 4B).

Pyocyanin (1 mg, 0.0048 mmol) was dissolved in DMSO-*d6* (500 µL) in a standard 5 mm NMR tube, and the spectrum was recorded. I-152_ox_ (2.52 mg, 0.0048 mmol) was then added, and the spectrum was recorded 20h after addition (Fig. S2).

### 2.7 PYO and I-152 Mass Spectrometry evaluation

Pyocyanin (PYO) and PYO with I-152 samples were resuspended at the above concentrations (paragraph 2.7) in PBS. The analyses were performed using Vanquish ultra-high-performance liquid chromatography (UHPLC) system, coupled with high-resolution mass spectrometry (HRMS) Orbitrap Exploris 240 (both supplied by Thermo Fisher Scientific, Waltham, MA). For targeted analysis, the mobile phases for chromatography separation consisted of LC-MS grade water (phase A) and LC-MS grade acetonitrile (phase B), both containing 0.1% of Formic Acid (all solvents supplied by Thermo Fisher Scientific, Waltham, MA). The chromatographic separation was performed with C18 Hypersyl GOLD column 150 × 2.1 mm, 1.9 μm (Thermo Fisher Scientific, Waltham, MA,) at 40°C and with a flow rate of 0.300 ml/min.. The gradient elution program was set as follows: 0-1 min, 0% B; 3 min, 40% B; 5.5-7.6 min, 90% B; 7.7-10 min, 0% B. The acquisition was performed in positive ion polarity mode with a calibration procedure conducted before the analysis; internal calibrants were also used in each run. The mass spectrometer was operated with the following parameters: for MS1 acquisition, the scan range was set to m/z 100–1000, with a resolution of 60,000 at m/z 200. The automatic gain control (AGC) target was set to 10e5, with automatic maximum injection time enabled. For MS2 acquisition, the m/z range was set to automatic, and stepped higher-energy collisional dissociation (HCD) was applied using normalized collision energies of 30%, 70%, 90%, and 150%. The resolution was set to 30,000 at m/z 200, with an AGC target of 10e4 and a maximum injection time of 54 ms. Identification and quantification of the targets were performed using FreeStyle 1.8 software (Thermo Fisher Scientific, Waltham, MA) and QuanBrowser software (Xcalibur 4.2, Thermo Fisher Scientific, Waltham, MA). For the untargeted analysis the samples were analyzed in both positive and negative ion polarity mode with calibration procedure performed before the analysis and with internal calibrants in each run. Both positive and negative chromatographic separations were performed using a reversed-phase C18 Hypersyl GOLD column (150 ×2.1 mm, 1.9 μm, Exploris 240 Thermo Fisher Scientific, Waltham, MA, USA). The mobile phases A for negative- and positive-mode analysis consisted of water with 10 mM ammonium acetate at pH 8 and water with 10 mM ammonium formate at pH 4.5, respectively, while acetonitrile with 0.1% formic acid was used as phase B for both analyses. The flow rate was 0.3 ml/min, with the column oven at 40 °C and the autosampler at 4 °C. The gradient elution program was set as follows: 0.8 min, 0% B; 20 min, 60% B; 24–28 min, 98% B; 28.2–32 min, 0% B. The mass spectrometer was operated with an MS1 range of 70–900 m/z, a resolution of 120,000 at m/z 200, an AGC target of 1 × 10[, and automatic maximum injection time. For MS2, an automatic m/z range was used, with stepped HCD normalized collision energies (20%, 50%, 80%, and 150%), a resolution of 30,000 at m/z 200, an AGC target of 2 × 10[, and a maximum injection time of 70 ms. The untargeted analysis was conducted using the AcquireX deep scan software (Thermo Fisher Scientific, Waltham, MA) with 5 ID runs, 3 replicates each sample and with PYO used as procedural blank. Data processing was performed using Compound Discoverer (Thermo Fisher Scientific, Waltham, MA), version 3.4.

### 2.8 Pyocyanin (PYO) quantitative assay in *P. aeruginosa* cultures

Treatment with I-152 (3 mM), colistin (4 mg/L), or both was added to the diluted overnight culture as described above. PYO production was determined at 72h as described by Shouman *et al.* (2023) [21], with slight modifications, by extracting the pigment from culture supernatants. The cells were removed by centrifugation at 6,000×g for 10 min at +4°C. Then, PYO in the supernatant was extracted by mixing 1 mL of supernatant with 600 µL of chloroform. PYO was then purified with 1 mL of acidified water (0.2 mol/L HCl), which gave a pink–red solution. For quantification of PYO in solution, absorbance was measured at 520 nm using a Tecan Spark® Multimode Microplate (Männedorf, Switzerland). Then the PYO concentration (µg/mL) was calculated as follows: OD520 × 17.072 [molar attenuation coefficient].

### 2.9 ROS evaluation in macrophages

The Total ROS-ID detection kit (Enzo Life Sciences, USA) was used according to the manufacturer’s protocols to evaluate the total ROS production in cells stimulated with PYO from *P. aeruginosa* and pre-incubated with I-152 (0.25 mM). RAW 264.7 cells (n=3 biological replicates and 3 technical replicates) were seeded with a density of 0.4 × 10^^6^ cells/well in a 6-well plate and were incubated with or without the compound for 2h before PYO (500 µM) addition. The level of ROS, namely hydrogen peroxide (H_2_O_2_), peroxynitrite (ONOO^-^), and hydroxy radical (•OH), was determined at the inverted microscope Axio VertA1 (Carl Zeiss, Germany) via GFP filter (Ex./Em. 470/525) whereas cell nuclei were stained with Hoechst and acquired via DAPI filter (Ex./Em. 352/454). Cells were subsequently split using 10% Trypsin, and the fluorescence signal was also evaluated via flow cytometry (Beckton Dickinson FACSscan), acquiring ∼10,000 cell events for each sample.

### 2.10 BMDM: isolation, culture, and treatments

Cells were obtained from 3 transgenic CFTR^-/-^ (B6.129P2-Cftrtm1Unc) (CF) and 3 CFTR^+/+^ (WT) C57BL/6 mice. After the sacrifice, bone marrow (BM) was collected. Briefly, bones were ground with a mortar and pestle, then filtered into a 50 mL vial containing PBS to remove any residual muscle tissue. The bone marrow was centrifuged at 100×g for 6 min; the pellet was resuspended in Dulbecco’s Modified Eagle Medium High Glucose (DMEM) supplemented with 10% heat-inactivated FBS, 2 mM L-glutamine, 100 U/mL penicillin, 100 μg/mL streptomycin (Gibco, Thermo Fisher Scientific, USA), 20 ng/mL recombinant macrophage colony-stimulating factor (M-CSF) (ConnStem Inc., USA), and put in culture in a 75 cm^2^ flask overnight. The next day, non-adherent cells were transferred to a 175 cm^2^ flask and allowed to differentiate for 7 days. At this point, the BMDM were detached, counted, and plated at a concentration of 1×10^^6^ cells/well in 6-well plates (Ø35 mm dishes) (Corning Incorporated, USA) for 24h before treatment. CF and WT cells were incubated with *P. aeruginosa* LPS (10 μg/mL) for 1h, after which I-152 (0.25 mM and 3 mM) was added to the media for 2h. The culture medium was then collected for cytokine release evaluation, and cells were lysed for gene expression analyses.

### 2.11 Real-time PCR

Total RNA was extracted using the RNeasy Plus mini kit (Qiagen, Germany) after lysing the cells with RLT buffer supplemented with 1% βmercaptoethanol (Sigma, Germany). According to the manufacturer’s instructions, cDNA was synthesized using Takara PrimeScript™ RT Master Mix (Takara, Japan) from 250 ng of total RNA. The RT-PCR reactions were performed on a QuantStudio 5 (Applied Biosystems, USA) in triplicate using PowerUp® SYBR Green Mastermix (Applied Biosystems, USA). The amplification conditions were: 40 cycles at 95°C for 10 min, 95°C for 10s, and 60°C for 50s. Relative mRNA expression was determined with the 2^-ΔΔCt^ method using 18S as a reference.

### 2.12 *In vivo* study

C57BL/6 wild-type (WT) mice were purchased from the Jackson Laboratory (Bar Harbor, ME, USA) and bred in the Yale University Animal Facility in pathogen-free ventilated cages. The animals, all male, were 8-weeks old and weighted around 22-25 g at the time of the experiment. All procedures were performed in compliance with the NIH (National Institutes of Health) Guide for the Care and Use of Laboratory Animals, ARRIVE guidelines and were approved by the Yale University Institutional Animal Care and Use Committee. In a first experiment, mice (15 in total) were treated intraperitoneally (i.p.) with I-152 (140 mg/Kg diluted in physiological solution) (6 mice) or received the vehicle (6 mice). After ½ and 1 hours, mice were sacrificed, and thiol species were determined in the lungs using a high-performance liquid chromatography (HPLC) procedure previously developed for thiol quantification in mouse organs [22]. Three mice were sacrificed at time 0h for basal values.

In a second experiment, 8 mice received three doses of *P. aeruginosa* LPS (L8643; Sigma, MO, USA) over 3 days. LPS (12.5 mg) was administered to the mice via nebulization using a Pulmo-Aide Compressor (Natallergy, Duluth, GA). A 5-mL aliquot of solution was nebulized over 15 minutes. Four mice received I-152 (140 mg/Kg diluted in physiological solution) i.p. on day 1, 1h before and 5h after LPS nebulization, on day 2 and 3, 1h before LPS. Mice were anesthetized on day 4, and 200 µl of blood was recovered by retro-orbital puncture. Bronchoalveolar lavage fluid (BALF) was collected using standard methods [23]. The BALF was first passed through a 100-μm cell strainer and then centrifuged at 1,200 rpm for 10 min. The supernatants were collected and stored in aliquots at −80 °C until analysis. The protein concentration of the BALF supernatant was determined by Bradford assay according to the manufacturer’s protocol (ThermoFischer Scientific). Cell pellets were resuspended in PBS, counted, and used for flow cytometry for differential cell counting. Lung tissues were used for histological analysis.

### 2.13 Flow cytometry staining and gating strategy

The cells isolated from BALF were washed and fixed with BD Cytofix (BD Biosciences, CA) and acquired on a BD LSR II using BD FACSDiva software. Data analyses were performed using FlowJo software (TreeStar, Ashland, OR). The following antibodies were used for flow cytometry and cell sorting: anti-CD45-BUV395 (BD Horizon), anti-CD38-FITC (BioLegend) anti-CD11b-PE-Cy7 (Invitrogen), anti-CD64-APC (BioLegend), anti-CD24-BV-421 (Invitrogen), anti-Ly6C-BV605 (BD Horizon), anti-CD11c-BV-805 (BD Horizon), anti-Ly6-G-AF-700 (BD Pharmingen), anti-Siglec-F-PerCP-Cy5.5 (BD Pharmingen), anti-MHC-II-APC/Cy7 (BioLegend), anti-CD4-BUV-737 (BD Pharmingen), anti-CD8-BV-711 (BioLegend), anti-B220-PE (BD Pharmingen) and LIVE/DEAD-BV-510 (BD Horizon). The gating strategy is adjusted from Misharin *et al.* [24] and is shown in Fig. S3. Single events were gated, followed by exclusion of dead cells stained with LIVE/DEAD. Small debris was excluded by the exclusion of FSC-H SSC-H small events, and from here, CD45^+^ cells were gated. As a next step, we gated alveolar macrophages (AMs) by first selecting for CD11c^+^ cells. From the CD11c^+^ fraction, we distinguished dendritic cells (DCs) from AMs by CD64/CD24 expression levels. Within the AM population, we distinguished monocyte-derived AMs (moAMs) as SiglecF^intermediate^CD38 ^high^ from tissue-resident AMs (trAMs) as SiglecF^high^ CD38 ^intermediate^. From the CD11c fraction, we gated on the remaining myeloid cells based on CD11b expression. Granulocytes were then gated from the CD11b^+^ cells based on high CD24 expression and low MHC-II expression. Granulocytes were then split into neutrophils as Ly6G^+^SiglecF and eosinophils as Ly6GSiglecF^+^. Lymphocytes were stained separately and identified as CD4^+^ CD4 T cells, CD8^+^ CD8 T cells, and B220^+^ B cells among singlet viable CD45^+^ cells.

### 2.14 Lung histological analysis

Mouse lung tissues were fixed in 10% neutral buffered formalin and paraffin-embedded according to routine methodology. Four µm-thick sections were stained with Hematoxylin and Eosin (H&E) (ST Infinity H&E Staining System, Leica Biosystem, Nussloch, Germany) for morphological analysis. Images were digitized using the microscope Eclipse Ci-L (Nikon Corporation, Tokyo, Japan) and NIS-Elements BR-2 software (v. 5.10; Nikon).

### 2.15 Magnetic multiplex immunoassay for profiling cytokines

Cytokines were measured in BMDM culture media, BALF, and serum. All samples were spun down at 10,000 g for 10 min at 4°C to remove cell debris, then stored at -80°C until analysis. The murine 8-plex kit was purchased from Bio-Rad (USA) and used according to the manufacturer’s recommendations. The kit detected the following cytokines: IFN-γ, IL-1β, IL-2, IL-6, IL-10, IL-12 (p70), and TNF-α. Plates were read on a BioPlex 200 instrument (Bio-Rad, USA), and cytokine concentrations (expressed as picograms per milliliter) were calculated using a standard curve and the manufacturer’s software (Bio-Plex manager software, v.6.1).

### 2.16 Statistics

Statistics and graphical data were analyzed using GraphPad Prism 11.0.0 (USA). Differences between experimental groups were assessed with one-way ANOVA (with Tukey’s *post hoc* correction for Figs. 1, 2, 4, 5, and 6) and the K-W test (with Dunn’s correction for Fig. 3). Results were considered statistically significant at *p* <[0.05. Quantitative image analyses were performed with ImageJ (USA).

**Fig. 5.**
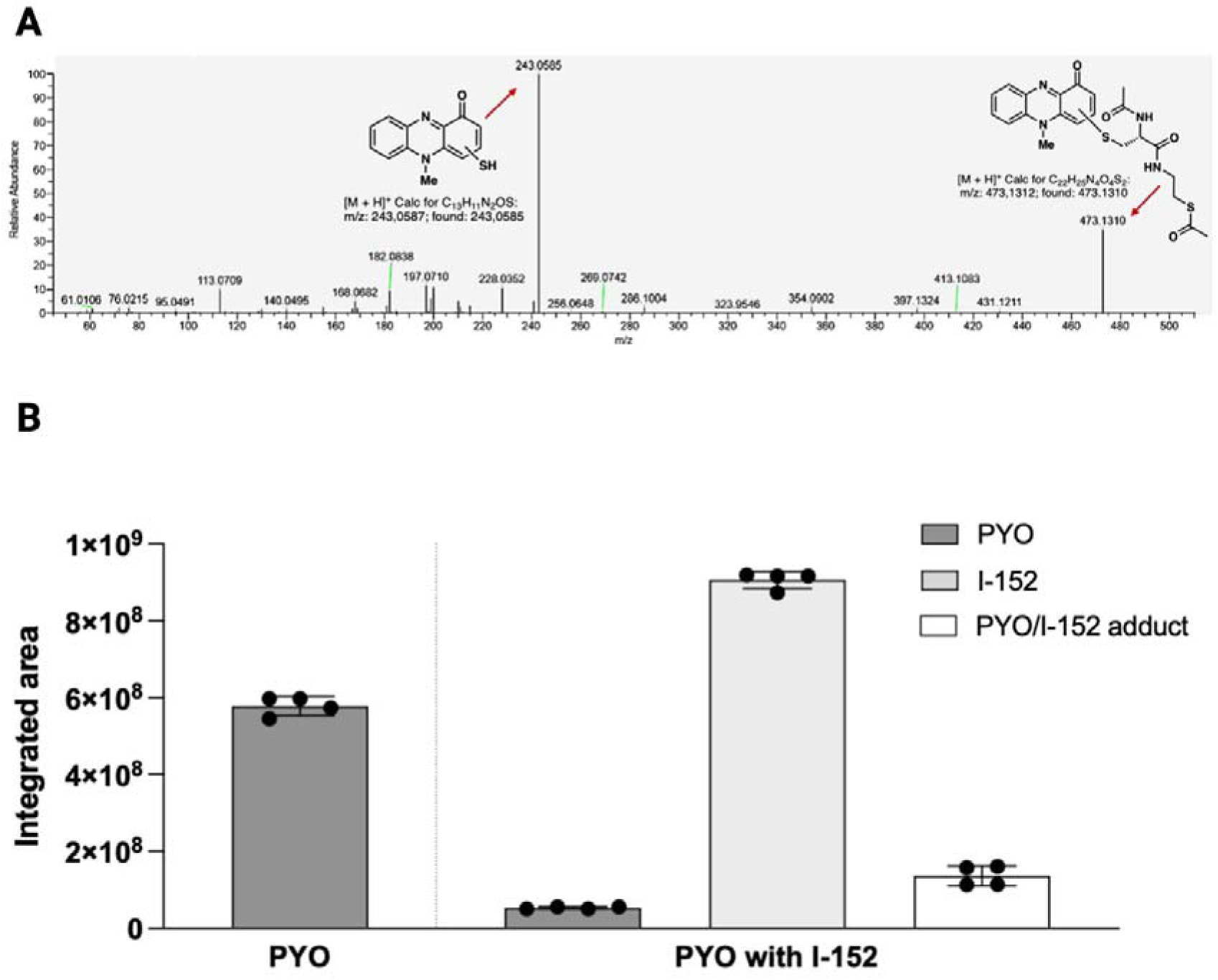
PYO levels after 96h of incubation in PBS. (**A**): MS/MS spectrum of the PYO/I-152 covalent adduct with m/z 473.1310 and assignment of the major fragments. (**B**): bar chart representing the peak areas of: PYO (dark gray) when incubated alone for 96h (5.66e08 counts/area) or in the presence of I-152 (5.3e07 counts/area), I-152 (light gray) (8.94e08 counts/area), and PYO/I-152 covalent adduct (white) (m/z 473.1310, 1.36e08 counts/area).

## 3. Results

### 3.1 I-152 effect on *P. aeruginosa* growth

I-152 is a conjugate of NAC and SMEA linked together by an amide bond. The molecule can cross the cell membrane, and metabolic studies on eukaryotic cells showed that it provides the dithiol derivative, which may play a crucial role in the release of NAC and MEA, two well-known pro-glutathione (GSH) compounds (Fig. S4) [14,19,25]. I-152 boosted GSH both *in vivo* and *in vitro* and showed bacteriostatic and bactericidal activity against *M. avium* [14]. I-152 toxicity was previously investigated [26–28], and recently, preliminary (Absorption, Distribution, Metabolism, and Excretion) ADME studies were conducted [29].

The antimicrobial activity of I-152 was evaluated by incubating *P. aeruginosa* in MHB in the presence of different I-152 concentrations (0.25, 1, and 3 mM). *P. aeruginosa* replication was compared with that of the bacteria grown in the absence of the molecule (CTR) at different times, *i.e.*, at 1h, 6h, and 24h. I-152 significantly interfered with *P. aeruginosa* replication in a concentration-dependent manner at 1h (Fig. 1A), but at subsequent time points, the bacterial load in treated samples was comparable to that in untreated samples (not shown). To ascertain the thiols that could be responsible for the antimicrobial effect registered at 1h, HPLC analysis aimed at identifying –SH carrying species was performed in untreated *P. aeruginosa* cells (CTR in Fig. 1B) or in 1 mM I-152-treated bacteria (I-152 in Fig. 1B). Only in the treated cells, I-152 was determined, while other thiol species, such as NAC, cysteine, or MEA, were not detected, and GSH levels were not increased with respect to the untreated, suggesting that the effect observed within the first hour may be ascribed to I-152 itself. Additionally, the bactericidal activity of I-152 was investigated by incubating *P. aeruginosa* in physiological solution for 1h at I-152 concentrations of 0.25 and 1 mM. The results displayed that low doses of the compound had a bactericidal effect on the pathogen at early time points (Fig. S5).

### 3.2 I-152 effect on planktonic bacteria growth and biofilm

Biofilms are complex communities of microorganisms enclosed in a self-produced extracellular matrix that helps them survive in unfavorable conditions. Bacteria inside the biofilm exhibit increased resistance against conventional antibiotics. In fact, the antimicrobial effect of these drugs is compromised because they cannot penetrate the biofilm matrix effectively [30]. On the other hand, misuse of antibiotics drives the evolution of traditional antibiotic resistance mechanisms in bacteria. Moreover, biofilm formation poses a challenge for host immune cells, favoring chronic infection [31]. For these reasons, nowadays, anti-pathogenic drugs that attenuate virulence or disrupt biofilms rather than killing bacteria are sought [32]. In our experimental model, following the schemes reported in Fig. 2, I-152 efficacy in inhibiting planktonic bacteria growth and biofilm formation was evaluated singularly and in combination with colistin. Thus far, this antibiotic is the only one effective in some cases of *P. aeruginosa* resistance, although higher rates of colistin resistance have recently been observed [33]. Two different experimental protocols were followed to determine whether the molecules could interfere with biofilm formation (Fig. 2A) or disrupt an established biofilm (Fig. 2D). Based on the results in Fig. 1, I-152 was administered at a concentration of 3 mM for 1h. Following the protocol shown in Fig. 2A, *P. aeruginosa* culture was treated with: I-152 for 1h before removal and subsequent addition of fresh medium for 23h (a), 4 mg/L colistin for 24h (b); the combination of 3 mM I-152 with 4 mg/L colistin for 24h (c). As reported in Fig. 2B, the analysis performed at 24h revealed that there were no effects of 3 mM I-152 in planktonic bacteria growth (a), while 4 mg/L colistin inhibited *P. aeruginosa* growth by about 90%, alone or in combination with I-152 (b and c). A reduction in biofilm formation (Fig. 2C) was provided by treatment a (about 44%) and by 4 mg/L colistin (about 83%), given either alone (b), or in combination (c). Then, we examined I-152 activity on the 18-hour-old pre-formed biofilm using the experimental scheme shown in Fig. 2D. This event resembles what occurs in several clinical situations, leading to significant morbidity, resistance to antimicrobial therapies, and immunological evasion [31]. In this experimental condition, the treatments were given after 18h of bacterial growth on the plastic surface of a multi-well plate. Once the biofilm was established, treatments were left for 6h. The results reported in Fig. 2E showed that I-152 (d) had no effect on the growth of *P. aeruginosa,* and that even colistin (e) had no discernible impact (about 11% inhibition), with significant improvement by I-152 addition (about 28% inhibition) (f). When examining the biofilm (Fig. 2F), colistin alone had no obvious eradication effect on mature biofilm (e), while the combination of the antibiotic with I-152 reduced its biomass by 25% (f).

For an in-depth study of the differences revealed by the most common biofilm biomass quantification technique, *i.e.,* the crystal violet (CV) biofilm assay, biofilm investigations were performed using atomic force microscopy (AFM). AFM is a high-resolution imaging technique that enables visualization of biofilm morphology and quantification of surface properties at the nanoscale level [34,35]. Biofilm physical factors, *e.g.,* elasticity and stiffness, may control adhesion and detachment, influencing *P. aeruginosa* virulence.

The results (Fig. 3) show the Force Modulation Microscopy (FMM) amplitude images of a formed *P. aeruginosa* biofilm after 18h (the required time for its formation), that was untreated (CTR), treated with 3 mM I-152 (I-152), or with 4 mg/L colistin (colistin), or with the combination of both (I-152+colistin) (Fig. 3A). The biofilm that received the combination was characterized by the overall decrement of FMM amplitude signal, matching with a reduced stiffness of the biofilm (Fig. 3B). Moreover, the FMM roughness parameters (Ra and Rq) revealed that the nanomechanical properties are heterogeneous along the biofilm surface, with regions particularly sensitive to the combined drugs, suggesting a patchy biofilm. These results suggest that I-152 can alter biofilm physical characteristics rather than bacterial growth and that, when combined with colistin, it can enhance drug penetration and antibiotic activity.

### 3.3 ^1^H NMR analysis and mass spectrometry evaluation of PYO and I-152

To further understand the I-152’s capacity to mediate *P. aeruginosa* biofilm matrix stabilization, we turned our attention to PYO, a redox-active phenazine whose activity can be significantly altered by redox cycling disruption, with a significant impact on biofilm formation [8,36]. Therefore, the inhibition of PYO production has been identified as an attractive antivirulence strategy for the treatment of *P. aeruginosa* infections [37]. To dissect whether I-152 could interfere with PYO itself, ^1^H NMR studies were performed. Initially, the redox properties of PYO were investigated using Hantzsch ester (HE) as a biomimetic hydride donor (Fig. 4A). We felt that Hantzsch esters might serve the role of small-molecule analogs of NAD(P)H, as their 1,4-dihydropyridine core has been shown to participate in hydride transfer to electrophilic π-systems in a variety of noncatalytic processes, including enone hydrogenation. Notably, this structural motif closely resembles the 1,4-dihydronicotinamide unit of NAD(P)H [38], which is well known for its electron- and hydride-donating properties and serves as the biological reductant of PYO [39].

A reaction between PYO and HE, in stoichiometric amounts, was carried out directly in an NMR tube in DMSO-*d_6_* and monitored over time by ¹H NMR spectroscopy in order to follow the evolution of the involved species. At the beginning of the reaction (Fig. 4Ac), three main species were identified in the spectrum: the Hantzsch ester (HE), the oxidized Hantzsch pyridine (HP), [40] and a third set of signals, upfield/shielded compared to PYO, that can be reasonably attributed to a reduced form of pyocyanin (PYO_red_). Notably, no additional signals characteristic of oxidized PYO were observed (expected at δ 8.13 – 7.74 and 6.27 – 6.11 ppm in the aromatic region and δ 3.91 for the methyl group), suggesting that reduction occurs rapidly under these conditions. After 20h (Fig. 4Ad), the Hantzsch ester signals disappeared, particularly the methylene resonance at δ 3.11 ppm, indicating its consumption. Simultaneously, an increase in signals corresponding to HP was detected, with characteristic aromatic resonances appearing in the region δ 8.52 ppm, consistent with the formation of the oxidized pyridine derivative. Over time (Fig. 4Ae), in the closed -tube NMR, the signal from the new species form of PYO_red_ disappeared.

Conversely, when the NMR DMSO solution of PYO and HE (Fig. 4Ad) was exposed to air, the (re)formation of PYO was observed (Fig. S1), thereby confirming that the newly formed species attributed to the reduced PYO was correctly assigned. The temporal correlation between HE consumption and HP formation, together with changes observed in the PYO signals, supports a hydride-transfer mechanism in which HE acts as a reducing agent. These spectral changes confirm that PYO undergoes reduction under the experimental conditions, highlighting its redox-active nature. This initial characterization provides a basis for understanding how I-152 may interfere with PYO-mediated processes, potentially by modulating its redox state.

A second experiment was performed using PYO and I-152 (Fig. 4B) under analogous conditions, with the reaction carried out directly in an NMR tube in DMSO-*d_6_*. The mixture was similarly monitored over time by ¹H NMR spectroscopy to evaluate whether I-152 can interact with PYO and alter its redox behavior. In this case, at the beginning of the reaction (Fig. 4Bi), four main species were identified in the spectrum: I-152, its oxidized form, a third species that could not be unambiguously assigned, and a fourth component displaying resonances consistent with those obtained employing the reduced agent HE and attributed to the reduced form of pyocyanin (PYO_red_). The presence of signals corresponding to oxidized I-152 alongside the most likely PYO_red_ suggests that a redox process may already be occurring at early stages of the experiment.

A third experiment was made from PYO and oxidized form of I-152 (I-152_ox_) (Fig. S2). Also, in this case, the reaction was carried out directly in an NMR tube using DMSO-d□. Both reagents PYO and the oxidized I-152 remain unaltered even after 20h at RT.

Overall, these observations support the hypothesis that I-152, through its thiol functional group, can interfere with PYO-mediated redox processes, potentially acting as a competing redox partner or modulating the equilibrium between PYO’s oxidized and reduced forms.

Moreover, it was observed that, when the reaction between PYO and I-152 was carried out in D□O or CD□OD (two solvents where both compounds were soluble), the species corresponding to PYO_red_ was not detected (data not shown). This behavior could be attributed to the intrinsic properties of DMSO, particularly its strong polarity and ability to act as a hydrogen-bond acceptor.

These characteristics may facilitate hydrogen-bond interactions with molecules, creating a microenvironment that can transiently stabilize the reduced form of PYO (PYO_red_). Such stabilization might alter the redox equilibrium and the kinetic profile of the system, potentially slowing reoxidation or favoring the persistence of the reduced species under the given experimental conditions.

Several attempts to isolate the reduced form of PYO (PYO_red_) were unsuccessful. In all cases, oxidized pyocyanin or degradation was observed.

To deepen the mechanisms by which I-152 can interfere with PYO, further studies were carried out in physiological solution. In the first instance, a targeted metabolomic approach was applied to samples containing PYO alone and to PYO incubated with I-152 for 96h in PBS to identify and quantify both molecules and to establish the influence of the thiol compound on PYO. HRMS data [m/z 473.1312 [M+H]^+^] and sulfur isotope pattern analysis showed the presence of compounds with a molecular formula of C_22_H_24_N_4_O_4_S_2,_ consistent with putative derivatives arising from S-conjugation of I-152 with PYO (most likely species that could not be assigned by ^1^H NMR) (Fig. 5A). In the same samples a marked decrease in PYO levels compared to the control (PYO alone) was observed (Fig. 5B). A difference of approximately one order of magnitude was observed in the integrated peak areas of PYO between the two samples; specifically, incubation with I-152 resulted in a significant reduction relative to PYO alone. Furthermore, the presence of potential adducts formed via the conjugation of I-152 with PYO was investigated based on previously reported phenazine/biothiol reactions [41]. However, a comparison of peak areas revealed a significant discrepancy; as shown in Fig. 5B, the sum of the residual PYO and the newly formed adduct accounts for approximately 30% of the starting material. This indicated that the formation of this adduct was insufficient to account for the total disappearance of PYO, suggesting the existence of additional unidentified species. To investigate this hypothesis and identify these further products, an untargeted metabolomics approach was adopted, enabling global profiling of reaction products, including those not initially anticipated. This profile contributed to the overall degradation of the starting material beyond the initially predicted outcomes. Among these, the most abundant species are reported in Table 1 provided in the Supplementary Material.

### 3.4 I-152 effect on PYO production in bacteria and PYO-mediated ROS generation in murine macrophages

In order to test whether PYO alterations by I-152 would be mirrored in the biological settings, PYO accumulation and PYO-induced ROS generation were evaluated. At 24h and 48h, PYO was not yet detectable, whereas its peak production, as indicated by the typical green-blue coloration, occurred at 72h, as previously reported [42]. As shown in Fig. 6A, only I-152, alone or in combination with colistin, led to a drastic decrease in PYO production, suggesting a possible correlation between *P. aeruginosa* biomass and PYO generation. Moreover, PYO promotes *P. aeruginosa*-related infection by inducing cellular oxidative damage [27,43]. PYO’s induction of oxidative stress is, at least in part, due to its ability to increase ROS intracellular levels and suppress antioxidant pathways. Macrophages participate in the first-line immune response against airway infections caused by *P. aeruginosa,* and we previously reported that I-152 activated the Nrf2-dependent antioxidant response and reduced *E. coli* LPS-induced ROS formation [27,43]. Based on these findings, we evaluated ROS formation in RAW 264.7 cells pre-treated with 0.25 mM I-152 for 2h and then stimulated with PYO for 30 min (Fig. 6B, C). Microscopic analysis performed with a kit detecting cellular ^•^OH, H_2_O_2_, and ONOO^-^ showed that the increase in ROS level was counteracted by I-152 pre-treatment (Fig. 6B). This result was confirmed by flow cytometry (Fig. 6C), further supporting the capacity of I-152 to interfere with PYO and/or counteract ROS formation.

**Fig. 6.**
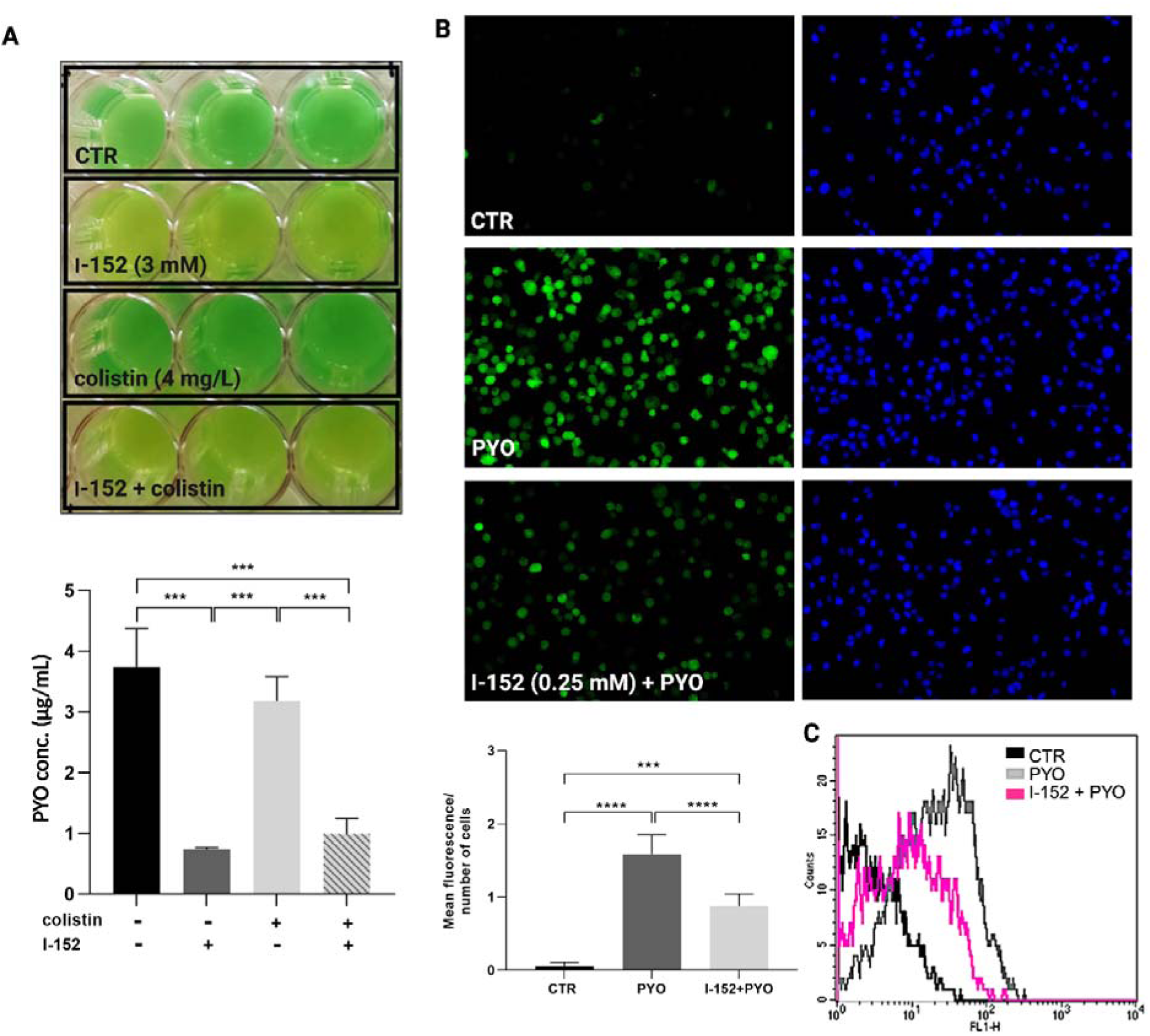
I-152 influence on PYO production and PYO-induced ROS. **(A**): PYO determination in the culture supernatants of *P. aeruginosa* without treatments (CTR), in the presence of I-152 (3 mM) and/or colistin (4 mg/L). Above: visual comparison of PYO release by *P. aeruginosa* in MHB at 72h; below: PYO quantification. PYO was extracted with chloroform and then re-extracted with 1 mL of acidified water (0.2 mol/L HCl). The absorbance was measured at 520 nm. B, C: PYO-induced ROS evaluation in RAW 264.7 cells. The reactive oxygen species (ROS), including ^•^OH, H_2_O_2_, and ONOO^-^, were detected using a ROS-ID^®^ Oxidative stress detection kit in: CTR (unstimulated), PYO (stimulated with 500 µM PYO for 30 min), and I-152 (pre-treated with 0.25 mM I-152 for 2h before PYO stimulation for 30 min). (**B**) Fluorescence microscope representative images of ROS (300 μm bar at 10× magnification); above: the intensity of green fluorescence corresponded to the amount of ROS quantified as the mean fluorescent intensity (green) normalized on cell nuclei stained with Hoechst 33342 (blue), below: bar graph depicting the mean fluorescence evaluation done using the ImageJ program. (**C**) Representative flow cytometry analyses of the aforementioned fluorescent cells for ROS detection. Data were expressed as mean ±LSD (n=3 biological replicates and 3 technical replicates). ***p<0.001; \*\*\*\**p*< 0.0001.

### 3.5 I-152 effect on inflammatory cytokine production in bone marrow monocyte-derived macrophages (BMDM) stimulated with LPS from *P. aeruginosa*

Although LPS is not a primary structural polymer of the matrix, it contributes to the biofilm’s electrostatic and surface properties through its highly charged outer membrane localization [30]. Similar to PYO, LPS triggers ROS generation and, in turn, macrophage activation via toll-like receptors (TLRs) [44]. LPS is a major component of the outer membrane in Gram-negative bacteria, and its interaction with macrophages leads to the production of inflammatory mediators, i.e., pro-inflammatory cytokines and ROS, that are pivotal for fighting the infection but could be harmful when overwhelming. CF macrophages exhibit a hyperinflammatory response when exposed to bacteria such as *P. aeruginosa* or its LPS [45].

To understand the impact of I-152 on cell response to LPS in WT and CF macrophages, inflammatory cytokine production, BMDM mRNA, and secretome were analyzed following a 3h *stimulus.* The investigated groups were: 1) CTR, a control group of BMDM which did not encounter any *stimuli*; 2) LPS, which received 1h LPS followed by 2h medium; 2-3) I-152 (0.25 mM or 3 mM), which received 1h LPS and then 2h of the respective treatment. The most secreted cytokines upon stimulation in the WT group were IL-6, IL-10, IL-12 (p70), and TNF-α (Fig. 7A). Concomitantly, the compound at the high concentration (3 mM) reduced all these upregulated cytokines, and even at the low concentration (0.25 mM), it exerted an effect against the main pro-inflammatory ones: IL-6 and TNF-α. In a similar trend, the mRNA levels of IL-6, IL-10, and IL-12 decreased with the high dose, whereas only IL-6 mRNA was abated with the low concentration (Fig. 7B). The TNF-α transcripts did not vary upon treatment, matching previous results described in the manuscript by Masini *et al.* [43]. CF BMDM showed a trend comparable to that of WT following LPS stimulation, with up-regulation of IL-6, IL-10, IL-12(p70), and TNF-α (Fig. 7C, D). The compound at both doses affected IL-6, IL-10, and TNF-α secretion (Fig. 7C); at the transcriptional level, IL-6, IL-10, and IL-12 were affected by the compound at the high dose, but only IL-6 was statistically downregulated by 0.25 mM I-152 (Fig. 7D).

**Fig. 7.**
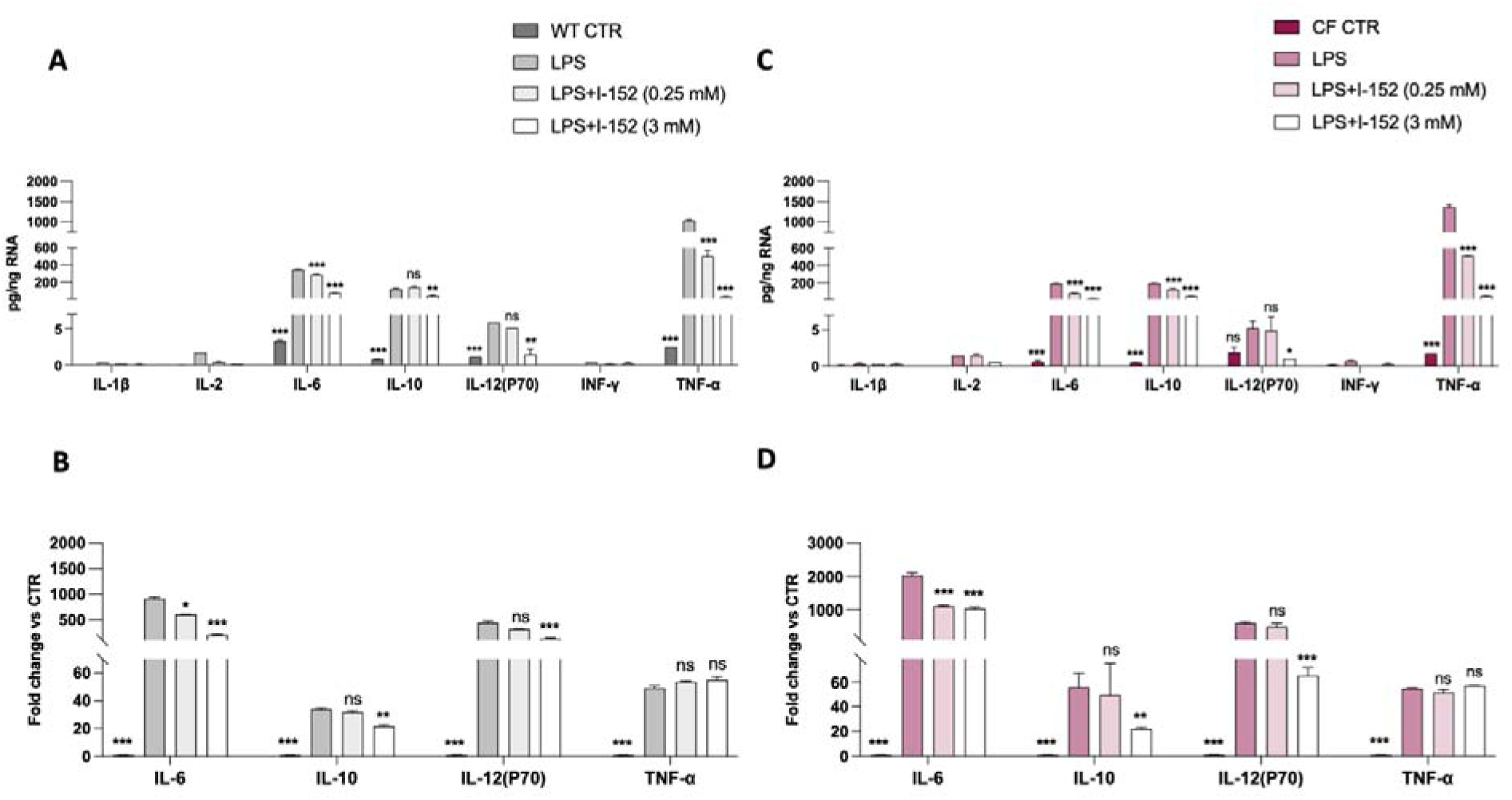
I-152 effects on LPS-induced cytokine expression in BMDM cells. The experimental groups were divided as follows: 1) CTR, a control group of BMDM which did not encounter any *stimuli*; 2) LPS, which received 1h LPS followed by 2 h medium; 3-4) I-152 (0.25 mM or 3 mM), which received 1h LPS followed by 2 h of the respective treatment. Bar graphs depict (**A**) WT mouse secreted cytokines, (**B**) WT mouse mRNA, (**C**) CF mouse secreted cytokines, and (**D**) CF mouse mRNA. mRNA levels were measured by quantitative reverse-transcription polymerase chain reaction (RT-qPCR) in total RNA extracted from BMDM. The 18S gene was used as a reference for relative quantification using the 2^-ΔΔCt^ method, and the results are expressed as fold change relative to the unstimulated CTR. Secreted cytokines were determined using a Bioplex array (pg/mL) and normalized to cell RNA content. The results represent the mean ± SD of 3 animals (3 CF and 3 WT). Statistical analysis was performed *vs* the LPS group. \**p*L<L0.05, **p<L0.01, ***p<L0.001.

### 3.6 I-152 effect on inflammatory response induced by *P. aeruginosa* LPS in mice

Acute lung infection due to *P. aeruginosa* is an increasingly serious problem that results in high mortality, especially in the compromised host. LPS can be recognized by the host immune receptor TLR4 and activate the innate immune response, leading to an anti-infection immune response [27,43]. Thus, we sought to examine the effects of I-152 in the context of acute pulmonary hyperinflammation induced by *P. aeruginosa* LPS exposure. Firstly, we evaluated whether systemic I-152 administration affected redox lung status. To this end, we investigated the thiol species content in the lung tissue at early times after intraperitoneal (i.p) I-152 administration (140 mg/Kg). This dosage, which has not been shown to cause adverse effects, was selected based on previous experiments in mice [26,28]. Actually, the ability of I-152 to deliver different thiol species, *e.g.*, NAC, MEA, and cysteine, and to increase basal intracellular GSH content in human and mouse cellular models, as well as in other mouse organs, has already been investigated elsewhere [27,28].

Lung thiols measured at 30 and 60 min after i.p. I-152 injection enabled precise quantification of the redox species derived from I-152 metabolism (Fig. 8). The results show that MEA and NAC reached peak levels (about 0.3 picomoles/µg organ) at 1/2h after I-152 administration, and then decreased at 1h (Fig. 8A). Cysteine levels were significantly higher in I-152-treated animals at both investigated time points (gray columns, Fig. 8B). The thiol species released by I-152 were conceivably used by the cells to make GSH, as suggested by the higher levels of the tripeptide measured at 1h (p=0.08) (Fig. 8A). Once established that I-152 could shift the redox balance of the lungs towards a more reduced state, we examined its anti-inflammatory effects in the next experiment using a *P. aeruginosa* LPS challenge (Fig. 9). On the first day, mice were treated i.p. with I-152 (140 mg/Kg) 1h before and 5h after LPS nebulization. Over the next three days, mice received I-152 1h before LPS nebulization (Fig. 9A). Mice stimulated with LPS and treated with the vehicle at the same times as those receiving I-152 were considered controls. Mice were sacrificed 24h after the last LPS nebulization, and bronchoalveolar lavage fluid (BALF) samples and lung tissues were collected (Fig. 9A). Moreover, serum was used to determine circulating cytokines. The complete blood count (CBC) did not differ between LPS- and LPS/I-152-treated mice, indicating that I-152 does not have a systemic negative impact on blood parameters (not shown). We assessed the immune cell profile in BALF samples from experimental mice using flow cytometry, following the flow strategy reported in Fig. S3 [46,47]. As expected, in response to LPS, the predominant cells recovered in the BALF samples were polymorphonuclear leukocyte neutrophils (PMN) (about 200×10^^4^ cells) and recruited monocyte-derived alveolar macrophages (moAM) (about 20×10^^4^ cells) (Fig. 9B). Mice treated with I-152 showed a trend toward reduced levels of PMNs, alveolar macrophages (AMs), eosinophils, and B cells, although these differences were not statistically significant, potentially leading to attenuated tissue damage as observed in Fig. 9C. No modulation of CD8^+^ and CD4^+^ T cells was observed among the experimental groups. By histology (Fig. 9C), in mice not receiving any stimulus (CTR), the bronchial and alveolar systems appeared normal. The terminal bronchioles are predominantly lined by non-ciliated Clara cells, which have distinctive apical cytoplasmic blebs (inset). The bronchiolar epithelial cells retain their ciliated appearance. The size and structure of the alveoli (air sacs) and the epithelial cells lining them remained unchanged. In the LPS-stimulated mice (LPS), a severe inflammatory reaction was observed, characterized by predominantly neutrophilic peribronchial (black arrow) and perivascular infiltration (red arrow) with alveolar exudation (macrophages/pneumocytes in the lumen) (asterisk). In the LPS+I-152 mice, there is clearly less alveolar exudation with no perivascular/peribronchial inflammation. Indeed, reduced alveolar membrane damage is also revealed by the lower protein content measured in the BALF of the treated mice (0.59±0.06 µg/mL in LPS mice vs 0.37±0.09 µg/mL in LPS+I-152 mice).

**Fig. 8.**
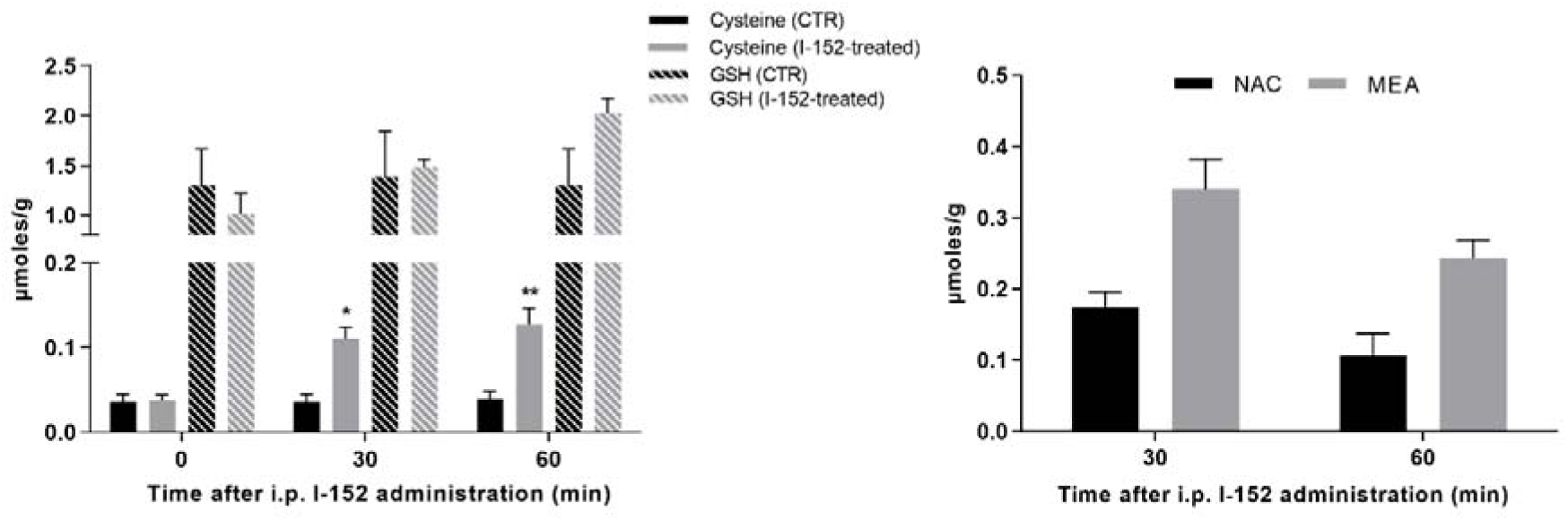
Lung thiol measurement. I-152 was administered i.p. at a concentration of 140 mg/Kg and after ½ and 1 hours, mice were sacrificed, and thiol species were determined through an HPLC method based on pre-column derivatization with DTNB that reacts with the –SH group [22]. Control animals received a placebo. The results represent the mean ± SD of 3 animals; *p<0.05, **p<0.01 vs CTR. Statistical analyses were conducted using an unpaired Student’s *t*-test, Welch corrected.

**Fig. 9.**
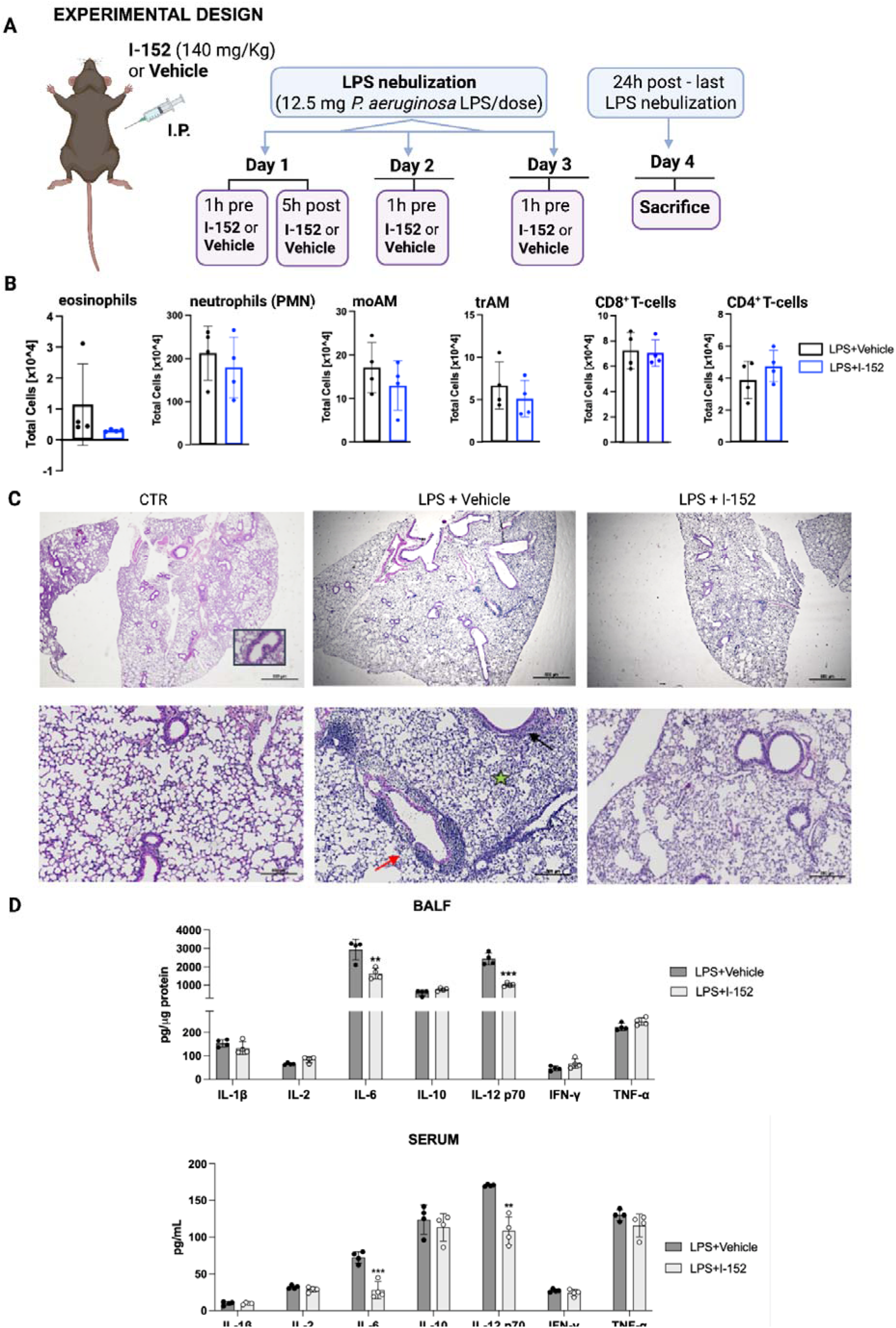
I-152 effects on inflammatory response in mouse lungs. (**A**): schematic cartoon showing the treatment regimen; mice received three doses of *P. aeruginosa* LPS over 3 days (one dose per day); on day 1, I-152 or the vehicle were administered 1h before and 5h after LPS nebulization; on the following days, the compound or the vehicle were administered 1h before LPS, and all analyses were performed 24h after the last LPS nebulization. (**B**): cell counts in the BALF. (**C**): representative hematoxylin-eosin staining of paraffin-embedded lung tissues. Magnification 2× (above) and 10× (below); scale bar: 500 µm; Inset: non-ciliated Clara cells. (**D**): Cytokine concentration in BALF and serum determined via Bioplex array. The results represent the mean ± SD of 4 animals; **p<0.01, ***p<0.001. Statistical analyses were conducted using one-way ANOVA (Tukey’s multiple comparison).

We further investigated the effects of I-152 on the hyperinflammatory response caused by LPS treatment by evaluating BALF and serum concentrations of the main immunomodulatory cytokines (Fig. 9D). Most of these cytokines are not detected in the BALF or serum of naive mice [46,47]. In the BALF of treated mice, we found lower levels of IL-6 and IL-12(p70), the cytokines most affected by LPSs, whereas the treatment did not affect the others. Regarding the circulating cytokines measured in serum, they mirrored the BALF trend, overall confirming the *in vitro* results obtained with BMDM (Fig. 7).

## 4. Discussion

*P. aeruginosa* is an opportunistic pathogen of immunocompromised and immunocompetent patients in whom wounds, burns, or foreign objects (i.e., catheters and ventilators) disrupt epithelial barriers, permitting the infection. Patients with an altered lung architecture, including those with cystic fibrosis (CF), are at particular risk for persistent *P. aeruginosa* colonization after an initial, acute infection. The intrinsic antimicrobial resistance of *P. aeruginosa* and virulence factors also contribute to its pathogenesis. In particular, biofilm plays a considerable role in persistence, colonization, multidrug resistance, and pathogenesis of *P. aeruginosa*. It is reported that 80% of chronic infections are associated with biofilms that form an obstinate physical barrier against antibiotics [48]. For this reason, future medical advancements will be based on the development of precise treatments that disrupt biofilm architecture. In this paper, we reported that a thiol-based molecule, named I-152, compromised the mechanical integrity of the biofilm, thereby potentiating colistin activity, which is known to be impaired by biofilm-embedded bacteria. In our experimental model, colistin alone was neither effective against mature biofilm nor against the initial-stage biofilm, whereas antibiotic activity was observed only when administered in combination with I-152.

Metabolism of I-152 in eukaryotic cells results in high levels of other thiol species, including NAC, which has been extensively investigated as an antibacterial and antibiofilm agent. In most studies [16,49,50], NAC has been shown to be effective in inhibiting biofilm formation and to act synergistically with antibiotics, but at higher concentrations compared to those used for I-152, even 20 times more, as also confirmed in this study (Fig. S6). Thiol quantification within *P. aeruginosa* cells revealed neither NAC nor other thiol species at early times of infection (Fig. 1B), suggesting that, at this time point, the observed effect may be ascribed to I-152 itself.

The observed biofilm perturbation was associated with I-152’s effects on PYO chemistry. PYO, being a redox-active blue phenazine pigment able to alter the redox equilibrium inside a biological system and with a relevant role in biofilm formation, was affected by the thiol molecule, leading to alterations in biofilm formation and resistance to antipseudomonal agents, *i.e.*, colistin [51]. The redox potential of PYO enables it to act as an electron shuttle, facilitating electron transfer between reducing agents (*e.g.,* NADH, NADPH) and molecular oxygen, thereby promoting ROS formation. I-152, thanks to its reducing capacity, can disrupt PYO redox cycling, favoring antibiotic activity in agreement with the results of Otero *et al.* [8]. PYO facilitates anoxic survival within established biofilms where cells experience oxygen limitation and redox stress [52]. By disrupting PYO-dependent electron shuttling to oxygen, I-152 could decrease PYO anoxic fitness remarkably at half the concentration of other reducing agents, *i.e*., DTT, TCEP [17]. Additionally, I-152 could act as a conjugating thiol partner for PYO, forming adducts likely contributing to reduce the PYO virulent effect, as previously postulated with NAC and GSH [53,54]. The performed untargeted analysis successfully identified several compounds, reflecting the complex interactions occurring within the system and highlighting a broad chemical diversity arising from concurrent events. These findings open up new perspectives on the reaction’s complexity, which may be addressed in future work. Other roles have been attributed to PYO, in particular as a pro-inflammatory agent [55], recently confirmed by Rodríguez-Urretavizcaya *et al.,* who showed that PYO led to a dose-dependent increase in pro-inflammatory cytokine levels that, however, were not normalized by an anti-PYO antibody that exhibited only cytoprotective effects on macrophages [56]. This finding suggests that PYO-induced ROS may serve as signaling molecules with a key role in the progression of the inflammatory process, and that this progression can be mitigated by quenching these ROS.

Overall, the decreased ROS level observed in macrophage cells exposed to a PYO insult and treated with I-152 could result from both PYO inhibition and ROS quenching, as already demonstrated in other systems [19,43].

Since PYO has been found in the sputum of CF patients chronically infected with *P. aeruginosa*, PYO likely contributes to *P. aeruginosa*-mediated pneumonia presumably through its ability to mediate damage through the generation of ROS, primarily superoxide (O_2_^•-^) and hydrogen peroxide (H_2_O_2_), and peroxynitrite (ONOO^-^) [57,58], resulting in DNA damage, disruption of membrane potential and redox balance, as well as activation of redox-sensitive signaling pathways that promote inflammation in the host [59]. A severe loss of GSH in airway epithelial cells exposed to PYO for prolonged time periods was also described [6]. GSH depletion has been attributed to the scavenging of PYO-derived ROS or to the direct oxidation of GSH by the toxin, depending on the experimental conditions [6]. By reducing ROS in immune cells (Fig. 4B), I-152 could decrease cytokine secretion in the host. Cytokine production is crucial for eliminating pathogens [60], however, the overproduction or dysregulation of these cytokines can lead to persistent inflammation, contributing to chronic inflammatory diseases such as CF and advanced stages of COPD [61]. Here, we demonstrated that the cytokine expression pattern in BMDM isolated from WT and CFTR-deficient mice under LPS stimulation can be balanced by I-152. We are aware of the complexity of CF pathophysiology, where excessive inflammation and the inability to resolve lung infections are the main hallmarks [62]. However, the important role of macrophages in contributing to the abnormal pro-inflammatory response in CF was previously highlighted [46]. The data show that in our experimental models, the most affected cytokines by LPS stimulus were IL-6, IL-10, IL-12, and TNF-α, and that I-152 inhibited IL-6, IL-10, and IL-12 expression both at transcriptional and protein levels in both WT and CF cells. This effect may be due to modulation of the activation of redox-sensitive transcription factors involved in the cytokine expression, i.e., NF-kB and AP-1, by I-152, as previously found in other models [43]. On the contrary, for TNF-α, whose expression is inhibited at the post-transcriptional level, I-152 could act at a later stage by interfering with ROS-mediated TACE activity up-regulation that is responsible for the conversion of inactive TNF-α precursor form to active mature form [62]. Regarding IL-10, this cytokine has been described as an anti-inflammatory defense mechanism developed by the immune system to control excessive production of pro-inflammatory molecules, limit tissue damage, and maintain or restore tissue homeostasis [63]. Indeed, an interplay and a mutual influence between IL-6 and IL-10 have been described in several pathologies, suggesting a complex regulatory mechanism. It is therefore difficult to predict the final outcome of I-152, as it will likely vary depending on the experimental conditions and levels of other cytokines [14].

These data suggest that I-152 could be a valuable tool for affecting *P. aeruginosa* pathogenesis in CF patients by targeting both the pathogen and the host. In fact, I-152 interferes with the production of virulence factors (i.e., PYO and biofilm) that enable colonization, pathogen survival, and the manipulation of the host’s defense mechanisms, thereby contributing to the pathogen’s success. In parallel, I-152 influences the host redox signaling network, which is implicated in the immune response. To confirm the *in vitro* anti-inflammatory effects of I-152, we conducted a preliminary *in vivo* experiment in WT mice stimulated with *P. aeruginosa* LPS. We observed attenuated cellular infiltration and significantly reduced IL-6 and IL-12 release both in the BALF and in the serum after 24h from the last LPS nebulization, while we did not find effects on TNF-α likely due to its kinetic; in fact, this cytokine is regulated at the translational level via the UA-rich sequence in the 3′ untranslated region of membrane undergoing proteolytic cleavage by a metalloprotease TNF-α-converting enzyme (TACE) as already discussed above [62]. Moreover, these results suggest that *in vivo* regulation kinetics may be mutually determined by complex molecular interactions and kinetic pathways that are difficult to fully recapitulate *in vitro* [64]. As confirmed histologically, LPS exposure induced acute lung injury characterized by a robust inflammatory response, with infiltration of neutrophils, macrophages, and other immune cells into the lung tissue (Fig. 9). This was accompanied by increased production of pro-inflammatory cytokines and chemokines, pulmonary edema due to increased alveolar-capillary permeability, and tissue injury, ultimately leading to hypoxemia. The beneficial effects of I-152 are likely due to changes in the redox state in the lungs, as evidenced by increased levels of NAC, MEA, cysteine and GSH after i.p. administration.

To fully appreciate the immune response and I-152 immunomodulatory activity, temporal pro-inflammatory cytokine patterns should be assessed *in vivo* at multiple time points. However, to limit animal distress, only one time point has been selected. Additionally, the small sample size (four mice per group) has been kept to a minimum in accordance with the ethical guidelines for animal research, which may limit the generalizability of the results. Despite these limitations, this study strongly suggests that I-152, whose absence of toxicity at the dose employed in this investigation had been previously observed [26,28], could counteract the exaggerated cellular and molecular responses in mice following nebulization with LPS from *P. aeruginosa*. Hence, I-152 treatment could be a therapeutic strategy able to efficiently hamper *P. aeruginosa* infection, acting against multiple targets, *i.e.*, dense biofilm and excessive inflammation that mutually reinforce each other, making infection control difficult and favoring drug resistance if only one aspect is targeted, as recently reported [65]. These promising results encourage future investigation in a chronic *P. aeruginosa* lung infection model that has been recently demonstrated as the most predictive mouse model for evaluating the efficacy of anti-inflammatory therapy [66] and/or in a CF mouse model to establish whether this compound could provide a promising solution for pulmonary infections caused by *P. aeruginosa*. We can conclude that I-152 may be considered a valuable candidate for the development of new multifunctional therapeutics to treat *P. aeruginosa* infections by interfering with the structure/function of virulence factors used by pathogens to colonize, invade, and persist in a susceptible host rather than killing the pathogen [67].

## Abbreviations

AFM: Atomic force microscopy
BALF: Bronchoalveolar lavage fluid
BMDM: Bone-marrow monocyte-derived macrophages
CF: Cystic fibrosis
CFTR: Cystic fibrosis transmembrane conductance regulator
CYS: Cysteine
GSH: Reduced glutathione IFN-γ Interferon-γ
IL: Interleukin
I.P: Intraperitoneal injection
LMW: Low molecular weight thiols
LPS: Lipopolysaccharide
MEA: β-mercaptoethylamine
MHB: Müller-Hinton broth
NAC: N-acetylcysteine
OD: Optical density
PBS: Phosphate-buffered saline
PYO: Pyocyanin
ROS: Reactive oxygen species
RT: room temperature
TNF-α: Tumor necrosis factor-α
WT: Wild-type

## Acknowledgments

The graphical abstract and Figs. 2, 9 were created using BioRender.com. The access to Biorender was supported by Dr. Juho Parvianen. Furthermore, we would like to thank POR MARCHE FESR 2014/2020. Asse 1, OS 2, Azione 2.1 - Intervento 2.1.1 – Sostegno allo sviluppo di una piattaforma di ricerca collaborativa negli ambiti della specializzazione intelligente. Thematic Area: “Medicina personalizzata, farmaci e nuovi approcci terapeutici”. Project acronym: Marche BioBank www.marchebiobank.it. «The content of the paper is the sole responsibility of the authors and can under no circumstances be regarded as reflecting the position of the European Union and/or Marche Region authorities».

## CRediT authorship contribution statement

**Michela Bruschi**: Conceptualization, Software, Investigation, Formal analysis, Data curation Writing - Original Draft, Visualization. **Sofia Masini**: Investigation, Visualization. **Francesco Palma**: Investigation, Visualization. **Yang Xiaoqiu**: Investigation, Data Curation. **Braga Cassia Lisboa**: Investigation, Data Curation. **Matteo Gregori**: Investigation, Visualization. **Cecilia Bucci**: Investigation, Data Curation. **Francesca Bartoccini**: Investigation, Methodology, Data Curation, Writing - Review & Editing. **Michele Menotta**: Investigation, Data Curation, Writing - Review & Editing. **Elisabetta Manuali**: Visualization, Writing - Review & Editing. **Liliana Minelli:** Investigation, Visualization. **Daniela Ligi**: Investigation. **Ferdinando Mannello**: Writing - Review & Editing. **Francesca Monittola**: Data curation. **Carolina Zara**: Investigation. **Caterina Di Pietro**: Investigation. **Rita Crinelli**: Review & Editing. **Giorgio Brandi**: Conceptualization, Methodology, Resources, Validation, Writing - Review & Editing. **Giovanni Piersanti:** Conceptualization, Methodology, Resources, Validation, Writing - Review & Editing, Fundings. **Emanuela Bruscia**: Conceptualization, Methodology, Resources, Validation, Writing - Review & Editing, Resources, Supervision. **Giuditta F. Schiavano**: Conceptualization, Methodology, Validation, Writing - Review & Editing, Resources, Supervision. **Alessandra Fraternale**: Conceptualization, Methodology, Validation, Writing - Review & Editing, Resources, Project administration, Funding acquisition.

