## Supplementary material for "An *N,S*-acetylated L-cysteine-cysteamine conjugate hinders pyocyanin redox cycling to weaken *Pseudomonas aeruginosa* biofilm and dampens LPS-driven acute pulmonary inflammation"

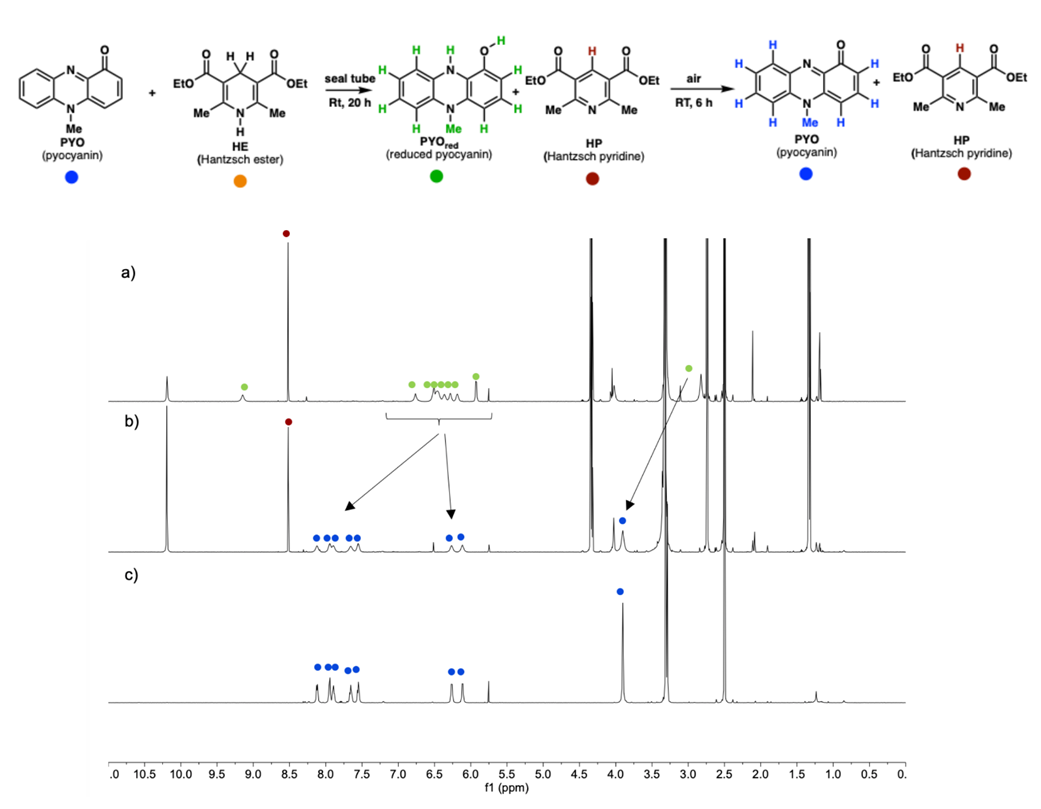

**Figure S1.** ^1^H NMR spectra in DMSO-d_6_ of (a) PYO (1 equiv) and HE (1 equiv), RT, 20 h; (b) air, RT, 6 h; (c) PYO.

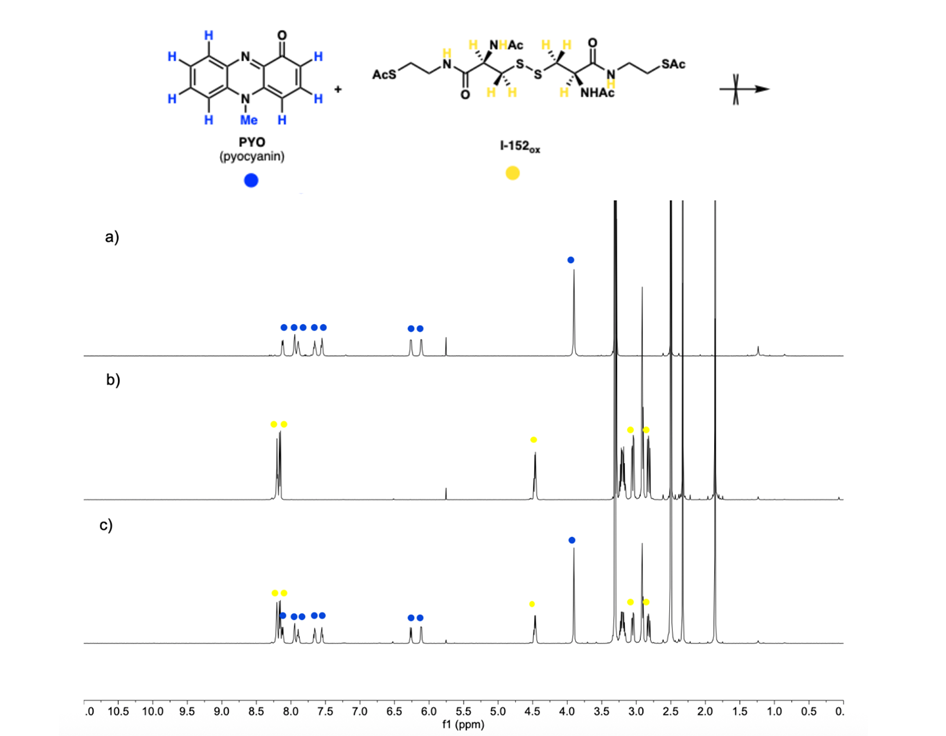

**Figure S2.** ^1^H NMR spectra in DMSO-*d_6_* of (a) PYO; (b) I-152_ox_; (c) PYO (1 equiv) and I-152_ox_ (1 equiv), 20 h, RT.

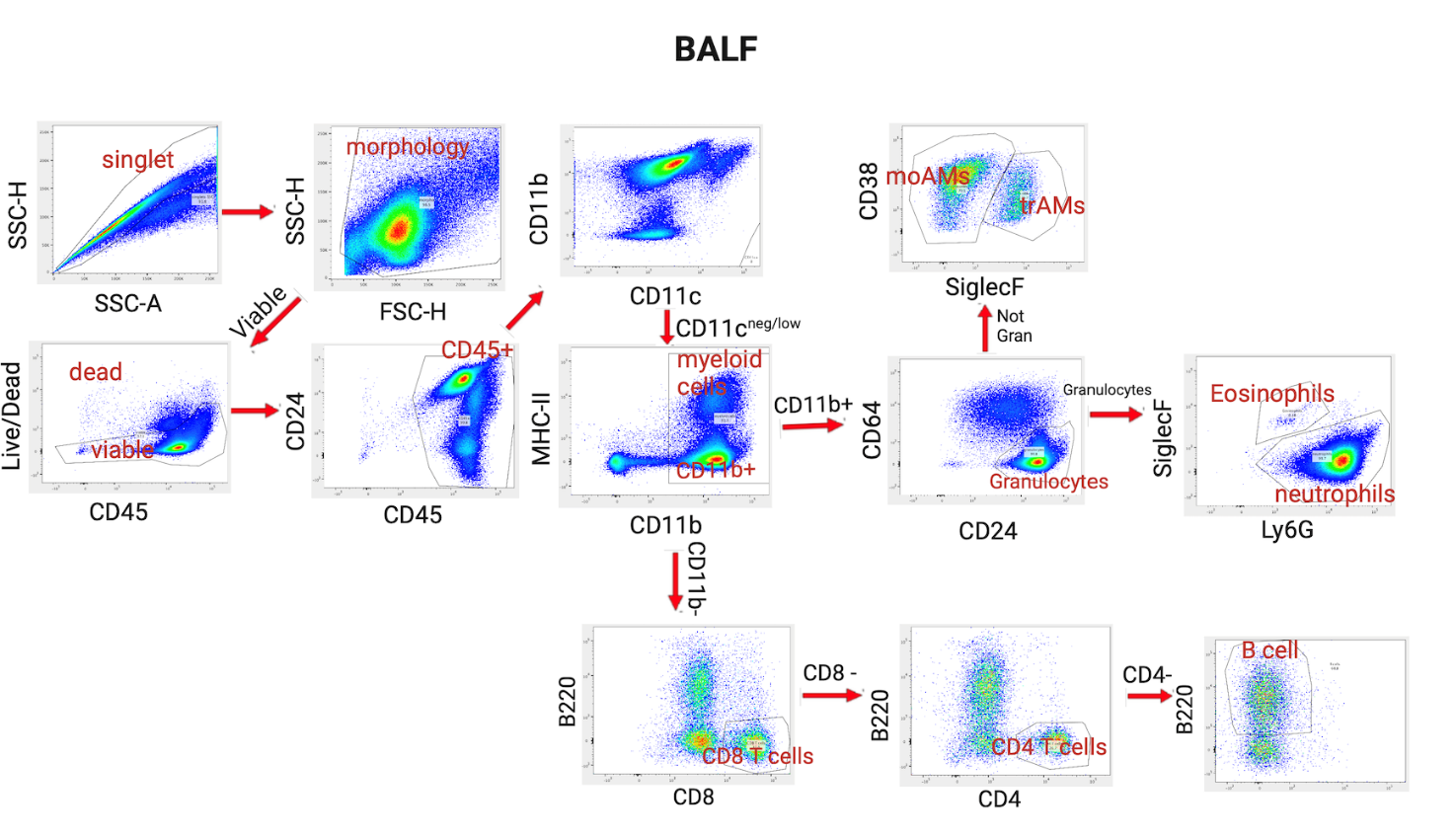

**Fig. S3**. Gating strategy on BALF for assessed immune cells.

Populations:

1.Eosinophils

2.Neutrophils

3.Monocyte-derived alveolar macrophages (moAM)

4.Tissue-resident macrophages (trAM)

5.CD8+ T cells

6.CD4+ T cells

7.B cells

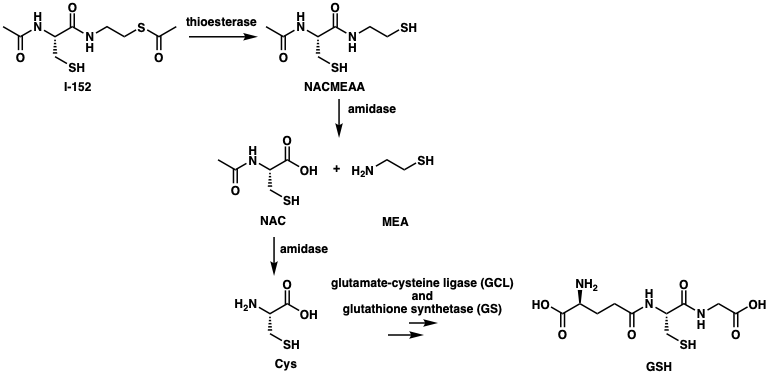

**Fig. S4**. Sketch of I-152 metabolism.

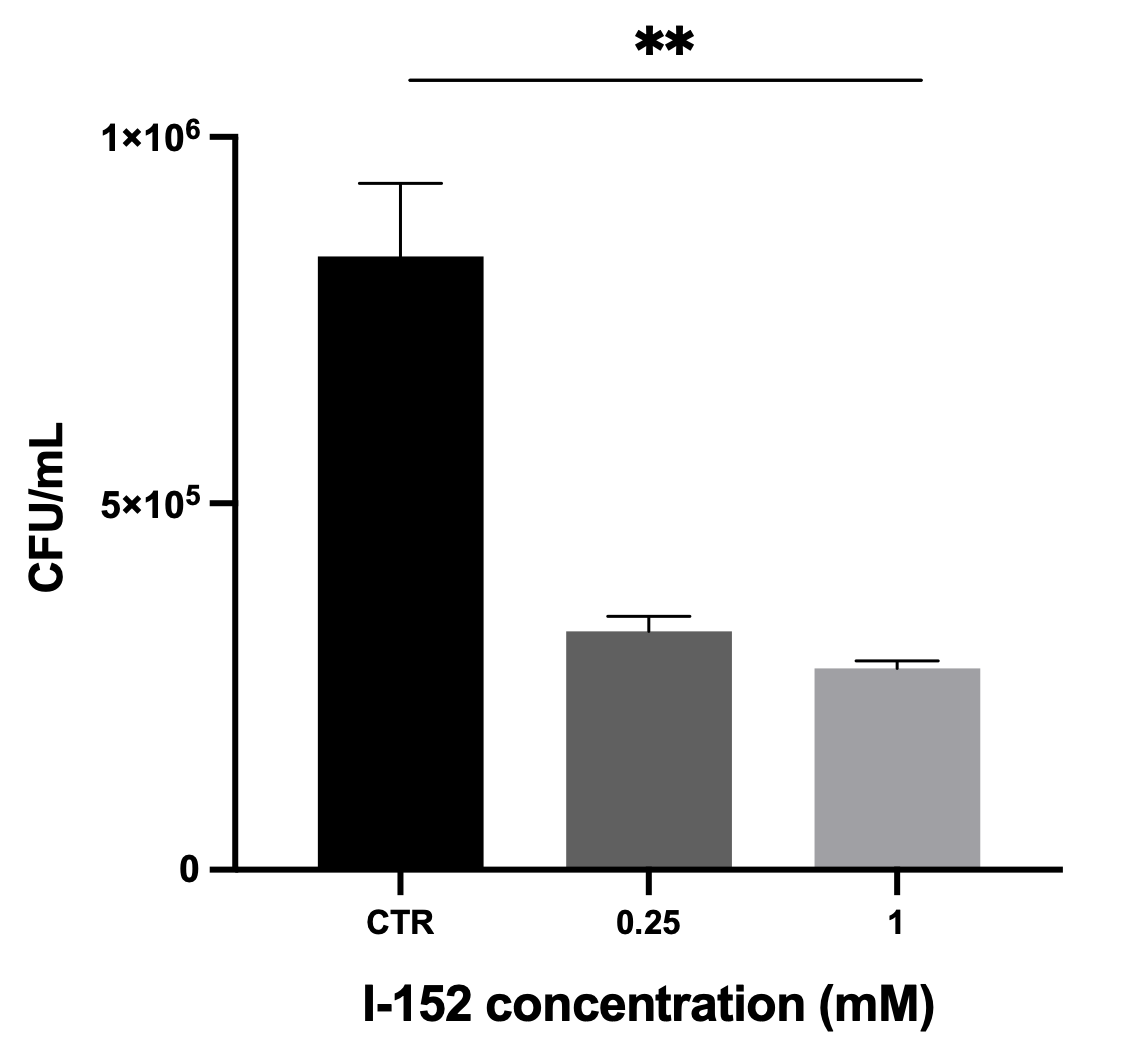

**Fig. S5**. I-152 bactericidal effect on *P. aeruginosa* evaluated in physiological solution at 1h.

| **Molecular formula** | **m/z value** | **Adduct** |
| --- | --- | --- |
| C₁₀HN₅O₂ | 222.0058 | [M-H]^-1^ |
| C₁₀H₂N₆ | 205.0269 | [M-H]^-1^ |
| C₁₀H₃N₅O | 208.0265 | [M-H]^-1^ |
| C₁₁H₁₈N₂O₄S₂ | 329.0599 | [M+Na]^+1^ |
| C₁₁H₂N₆ | 217.0268 | [M-H]^-1^ |
| C₁₁H₂N₆O | 233.0217 | [M-H]^-1^ |
| C₁₂H₂₁O₇P | 309.1094 | [M+H]^+1^ |
| C₁₄H₂₄N₄O₄S₄ | 485.0664 | [M+FA-H]^-1^ |
| C₁₄H₆N₂ | 102.0338 | [M+2H]^+2^ |
| C₁₆H₃₂O₂ | 274.2738 | [M+NH_4_]^+1^ |
| C₁₆H₈N₇O₃P | 376.0337 | [M-H]^-1^ |
| C₁₇H₃₁N₆O₃PS₄ | 571.1035 | [M+FA-H]^-1^ |
| C₁₉H₂₆N₈O₂S₄ | 527.1110 | [M+H]^+1^ |
| C₁₉H₂₇N₆O₄P₃S | 573.0986 | [M+FA-H]^-1^ |
| C₂₀H₄₆N₆O₄ | 452.3942 | [M+NH_4_]^+1^ |
| C₂₁H₂₂N₈O₃P₂S | 529.1066 | [M+H]^+1^ |
| C₂₂H₄₆O₅ | 408.3682 | [M+NH_4_]^+1^ |
| C₂₃H₄₈O₆ | 438.3786 | [M+NH_4_]^+1^ |
| C₂₆H₄₈N₄O₃ | 482.4045 | [M+NH_4_]^+1^ |
| C₂₆H₅₄O₈ | 512.4151 | [M+NH_4_]^+1^ |
| C₂₇H₅₀N₄O₃ | 496.4202 | [M+NH_4_]^+1^ |
| C₂₈H₅₂N₄O₄ | 526.4305 | [M+NH_4_]^+1^ |
| C₂₉H₃₄N₁₀O₄S₅ | 747.1452 | [M+H]^+1^ |
| C₂₉H₅₄N₄O₅ | 556.4412 | [M+NH_4_]^+1^ |
| C₃₀H₆₂O₁₀ | 600.4673 | [M+NH_4_]^+1^ |
| C₃₁H₄₈ClN₈O₁₇PS₃ | 484.0934 | [M+2H]^+2^ |
| C₃₁H₆₄O₁₀ | 614.4831 | [M+NH_4_]^+1^ |
| C₃₃H₆₂N₄O₇ | 644.4936 | [M+NH_4_]^+1^ |
| C₄H₉N₃P₂ | 81.5205 | [M+2H]^+2^ |
| C₄₂H₆₆NO₄P | 680.4797 | [M+H]^+1^ |
| C₄₄H₇₀NO₄P | 708.5108 | [M+H]^+1^ |
| C₆H₁₁NO₂S | 162.0582 | [M+H]^+1^ |
| C₆H₃N₃O₃ | 164.0102 | [M-H]^-1^ |
| C₇H₁₂N₂O₂S₂ | 265.0322 | [M+FA-H]^-1^ |
| C₇H₁₃NO | 128.1069 | [M+H]^+1^ |
| C₇H₁₅ClN₂O₃S₂ | 255.0035 | [M-H-H_2_O]^-1^ |
| C₇H₅N₆OP | 111.0202 | [M+2H]^+2^ |
| C₉HN₃O₄ | 213.9895 | [M-H]^-1^ |

**Table 1.** Overview of the major molecular species identified through untargeted metabolomics. The table provides a summary of the most significant ions (m/z) observed in both positive and negative ionization modes, including their assigned molecular formulas and the specific adduct forms identified. These chemical features were detected after 96h of incubation in PBS, and were selected based on their high relative abundance (cut off: 3×10^8^ counts/area).

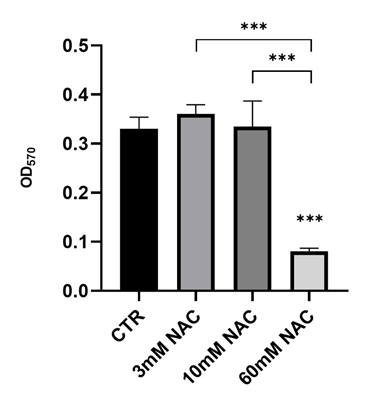

**Figure S6.** NAC effect on *P. aeruginosa* biofilm formation at 24h.

The biofilm biomass was stained with crystal violet.
